# Short-term sensory memory mediates adaptation, habituation, and a paradoxical neural-behavioral transformation in *C. elegans*

**DOI:** 10.1101/2025.08.08.669341

**Authors:** Hamilton White, Sruti Mallik, ShiNung Ching, Dirk R. Albrecht

**Affiliations:** Worcester Polytechnic Institute, Department of Biomedical Engineering; UMass Chan Medical School; Washington University in St. Louis, Department of Electrical and Systems Engineering; Worcester Polytechnic Institute, Department of Biology and Biotechnology; Worcester Polytechnic Institute, Department of Electrical and Computer Engineering

## Abstract

Repeated exposure to stimuli elicits decreasing sensory neural responses over time (adaptation). However, resulting behavioral responses can either weaken over time (habituation) or remain invariant, indicating that the neural-behavioral link is not fixed. To investigate neural adaptation and its flexible translation into behavioral decision making, we created a mathematical framework hypothesizing (1) that sensory networks optimize the speed and accuracy of encoding exogenous stimuli, and (2) that representations form along two time scales, one embedding immediate information and the other stimulus history. Using experimental recordings of the nematode *C. elegans*, we validated normative model predictions of this optimal encoding strategy, specifically how neural dynamics and adaptation levels vary with stimulus timing. A parametric Bayesian decoder architecture predicted conditions leading to behavioral habituation or invariance, but also paradoxical inversion, whereby appetitive stimuli elicit aversive responses. Experiments with food odors validated that inversion behavior occurred after several repetitions with a long stimulation time and low odor concentrations. Mechanistically, during sensory neural adaptation, weaker immediate stimulus representations can be compensated by secondary processes through memory effects, with biological origins that remain to be studied.

## Introduction

Behavioral habituation is a fundamental form of non-associative learning in which repeated exposure to the same stimulus leads to a decrement in behavioral response while preserving sensitivity to novel stimuli ***Groves and Thompson (1970***); ***Rankin et al. (2009***). This behavioral phenomenon is often attributed to neural adaptation, the progressive reduction in sensory neuron activity during repeated stimulation ***Pellegrino et al. (2017***). Although this view implies a direct link between neural adaptation and behavioral habituation, growing evidence suggests that the relationship is more complex ***Khan et al. (2022***). Here we explore olfactory adaptation in *Caenorhabditis elegans* using an integrated experimental and computational approach, identifying a pivotal role for transient neural representations of recent stimuli in shaping behavioral decisions. Our mathematical framework reveals that sensory systems construct representations on two timescales: an immediate component reflecting current stimulus features, and a slower component that integrates stimulus history. These dynamics allow for flexible behavioral outcomes that include not only habituation, but also response invariance and even paradoxical behavioral inversion, in which repeated appetitive stimuli can evoke aversive responses.

Sensory-to-behavior transitions in animals arise from robust, hierarchical neural architectures that are evolutionarily conserved across phyla ***Strausfeld and Hildebrand (1999***). In such systems, peripheral chemosensory receptor inputs are transduced into electrical signals that project through afferent pathways to the brain, where they are integrated and transformed before descending via efferent pathways to drive motor behaviors. This organization motivates our use of abstraction and computational modeling to examine (i) how latent representations of stimuli are dynamically constructed from early sensory activity, and (ii) how behavior results from the interplay between current sensory information and prior experience.

Our computational model attempts to capture key high-level aspects of hierarchical signal routing and transformation in the brain, within a computational paradigm that predicts both sensory neural activity and behavioral outputs. It builds upon normative theories of neural coding that assume the brain seeks to form internal representations that are accurate, energetically efficient, and robust to noise ***Ganguli and Simoncelli (2014***); ***Wei and Stocker (2015***). In this efficient coding formulation, sensory neural activity aims to: (1) **minimize inaccuracy** of the stimulus representation, (2) **minimize energy** costs associated with neural encoding, and (3) **minimize noise** or rapid fluctuations that may degrade information reliability.

This work extends a rich body of experimental and theoretical research on habituation, adaptation, and memory in invertebrate models such as *Caenorhabditis elegans* and *Drosophila melanogaster* ***Rankin et al. (1990***); ***Rankin and Broster (1992***); ***Beck and Rankin (1993***); ***Giles and Rankin (2009***); ***Linster et al. (2009***); ***Das et al. (2011***). For validation, we use *C. elegans* as the animal model due to its experimental tractability, evolutionarily conserved neurochemical pathways, compact and fully mapped 302-neuron connectome ***Hobert (2018***); ***White et al. (1986***); ***Cook et al. (2019***), and the availability of well-established experimental tools for noninvasive recording of *in vivo* neural activity ***Larsch et al. (2013***); ***Reilly et al. (2017***); ***Han et al. (2021***); ***White et al. (2023***) and quantification of sensorimotor behaviors ***Albrecht and Bargmann (2011***). Neural models of *C. elegans* have successfully reproduced several behavioral phenomena, including touch habituation ***Varshney et al. (2011***); ***Atanas et al. (2023***); ***Cangelosi and Parisi (1997***). However, while the *C. elegans* neural circuit is currently the most completely described, key elements remain incomplete, including the valence and dynamic regulation of synapses and extrasynaptic communication networks mediated by neuropeptides and other neuromodulatory signals ***Bargmann (2012***); ***Ripoll-Sánchez et al. (2021***); ***Watteyne et al. (2024***). Hence, a recent focus has been on understanding the algorithmic basis of habituation at the level of general principles that regulate its function, abstracting away specific details at the cellular and membrane level ***Marsland (2009***); ***Bargmann (2012***); ***Shen et al. (2020***). Our contribution builds on this top-down approach, proposing that adaptation and habituation emerge as outcomes of optimal strategies for stimulus representation under constraints.

### The central contributions of this work include

- **A normative modeling framework** that explains sensory adaptation as mathematical optimization of neural encoding under repeated stimulation, validated using recordings in *C. elegans*.
- **Generalization to novel conditions**, where the model is trained on a single stimulus pattern but it successfully predicts neural and behavioral responses under different temporal and intensity parameters.
- **Identification of a sensory memory mechanism** that mediates flexible behavioral outputs, including paradoxical inversion, where past exposure alters the valence of representations to repeated stimuli.
- **Validation of model predictions** in prospective experiments using repeated food odors pre-sented at different concentrations and durations.

These findings suggest that short-term sensory memory, shaped by stimulus history, allows the nervous system to flexibly reconfigure the mapping between stimulus and behavioral response, potentially serving as a general principle across sensory modalities and organisms.

## Results

### Behavioral invariance: repeated stimulation can cause neuronal adaptation without behavioral habituation

*C. elegans* respond to stimulation with probabilistic behavioral responses that enable locomotion toward favorable environments and away from harm. For example, appetitive (attractive) responses cause animals to move forward up a concentration gradient, whereas aversive responses include pauses, short (<1 s) reversals, and reorienting responses termed pirouettes, comprised of a long reversal and a nearly 180^°^ “omega” turn ***Gray et al. (2005***). Whereas individual responses to repeated stimulation are not predictable, the probability of each behavioral response is consistent across animal populations and repeats in an intensity-dependent manner ***Albrecht and Bargmann (2011***). Some stimuli respond reliably for hours (such as olfactory food cues), whereas others habituate quickly (e.g., gentle touch) ***Rankin and Broster (1992***).

Sensory neural responses to repeated stimulation can be visualized as increased fluorescence of the intracellular calcium reporter GCaMP imaged by optical microscope ***Larsch et al. (2013***). Neural adaptation is represented by decreasing calcium response magnitude and decreased fluorescence each trial. For example, repeated presentation of 20-s pulses of the food odorant diacetyl (0.4 *µ*M) in a microfluidic device (Figure 1A), once per minute, elicits robust calcium responses in the AWA chemosensory neurons that decrease in magnitude seven-fold over 24 trials (Figure 1B-D). Behaviorally, these attractive odor pulses rapidly increase forward locomotion probability, whereas stimulus removal increases pauses and reversal probability (Figure 1E-G). However, under these conditions, no behavioral habituation is observed, and behavioral responses remain roughly equivalent in magnitude across 24 repeated trials (Figure 1H). We term these consistent behavioral responses, despite neural adaptation, **behavioral invariance**.

**Figure 1.**
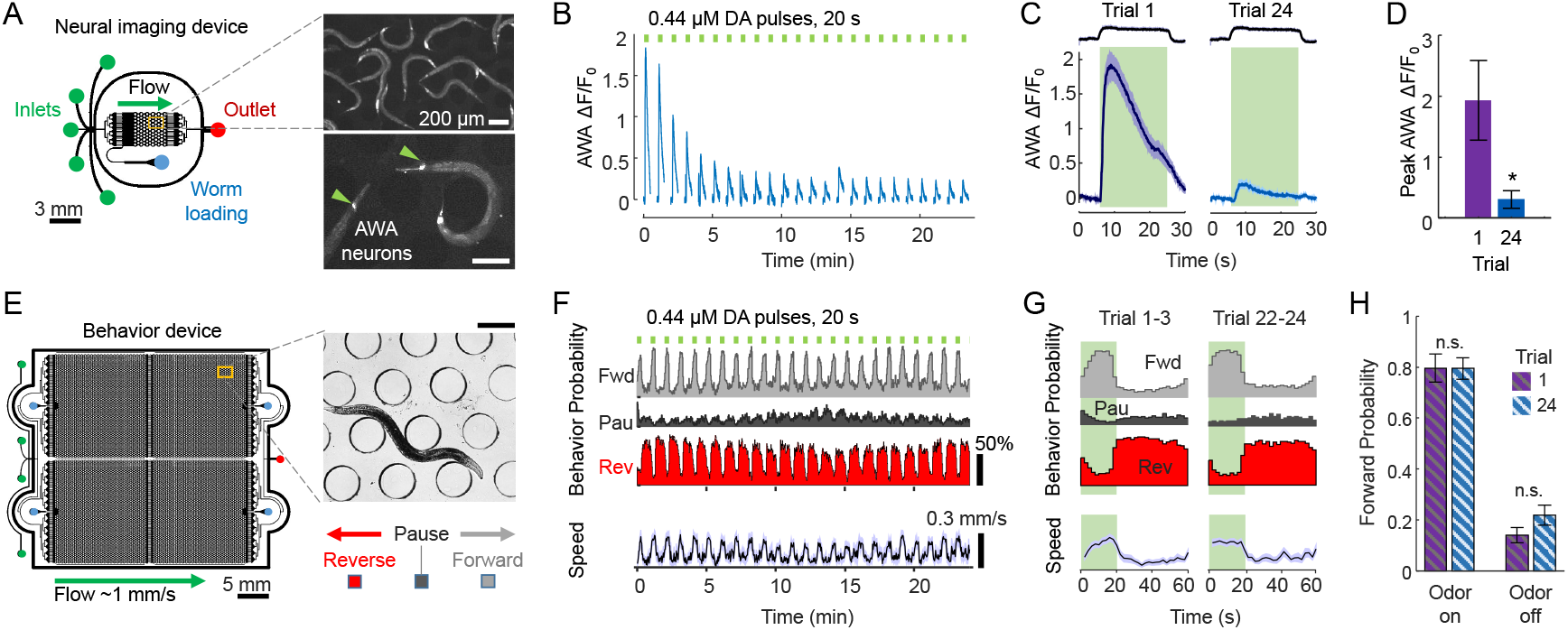
Repeated odor stimulation causes neural adaptation but invariant behavior responses. **A**. Microfluidic system for imaging neural responses to precisely-timed odor pulses. Inlets of a microfluidic device are switched to deliver stimulus or buffer to animals expressing the calcium sensor GCaMP in specific neurons, such as the AWA chemosensory neurons (inset arrowheads). Scale bars, 200 *µ*m. **B**. AWA neural activity adapts during 24 repeated pulses of diacetyl (DA) odor, 20s duration every 60s, indicated by mean relative fluorescence change (n = 13). **C**. Comparison of naive (trial 1) and adapted (trial 24) neural responses. Curves and shading represent mean and SEM, n = 13. **D**. Peak neural activity adapts over 6-fold from trial 1 to 24. Bars indicate mean ± SD, n = 13 animals. ^*^p = 0.0365. **E**. Behavior devices house ^∼^100 animals and record locomotory responses to stimulus pulses, including instantaneous behavioral states Forward, Pause, and Reverse (inset, scale bar 200 *µ*m). Forward behavior indicates animal preference for current conditions. **F**. Behavioral responses to 24 repeated pulses of odor as in panel B. Probabilities of Forward (Fwd) locomotion increase during odor presentation, whereas Pause (Pau) and Reverse (Rev) states decrease in odor. Locomotory speed also increases in odor. Data represent mean ± SEM shading, n = 28, binned in 2s increments. **G**. Behavior probabilities and speed, averaged for early (trials 1-3) and late (trials 22-24), indicate consistent (invariant) behavioral response dynamics over the 60s trial interval. Shading represents odor stimulation. **H**. Forward probability during odor presentation (left) or absence (right) is unchanged between early and late trials, indicating no habituation occurs under these stimulation conditions. Bars indicate mean ± SD, n = 28 animals. n.s., not significant.

### A dynamical model with latent sensory memory realizes neuronal adaptation and behavioral invariance

To investigate the factors that might lead to sensory neural adaptation, we developed a top-down mathematical model that realizes these trends. The computational model is based on the theoretical premise that sensory neural dynamics should extract meaningful information from environmental cues to aid higher level processes. Based on this premise, in ***Mallik et al. (2020a***), we introduced a normative model that enacted detection of solitary sensory cues on a single time scale. We posited that early olfactory networks create a dynamic representation (i.e., a neural code) in an energy efficient manner when a stimulus is presented and subsequently withdrawn. However, when an organism encounters the same stimulus repetitively, it would be seemingly wasteful for sensory networks to reproduce the same representation over and over again. Presumably, animals can leverage representations persisting over longer dynamic timescales (i.e., memory) to maintain sensory detection over longer temporal horizons. Based on this notion, we postulate that latent sensory representations are constructed and processed along multiple timescales, embedding a mathematical equivalent of short-term sensory memory. This latent information is passed on for higher-level processing which eventually generates behavior. We also provide hypotheses regarding how the preference of an organism to certain stimulus intensities might be integrated into the model. The model consists of three key components, presented thus:

#### Model part I: Optimization framework for dynamic sensory encoding

Figure 2 relates the flow of sensory information to behavior in the *C. elegans* neural circuit and the computational model. Note the close correspondence between experiment and model diagrams at the level of stimulus input, sensory neural response, and behavioral output (see also Box 1). Stimulus input patterns are precisely controlled in time and intensity using microfluidic systems, with sensory neural activity recorded by optical microscopy and fluorescent reporters, and behavior quantified by machine vision ***Larsch et al. (2013***); ***Albrecht and Bargmann (2011***); ***Lagoy (2018***).

**Figure 2.**
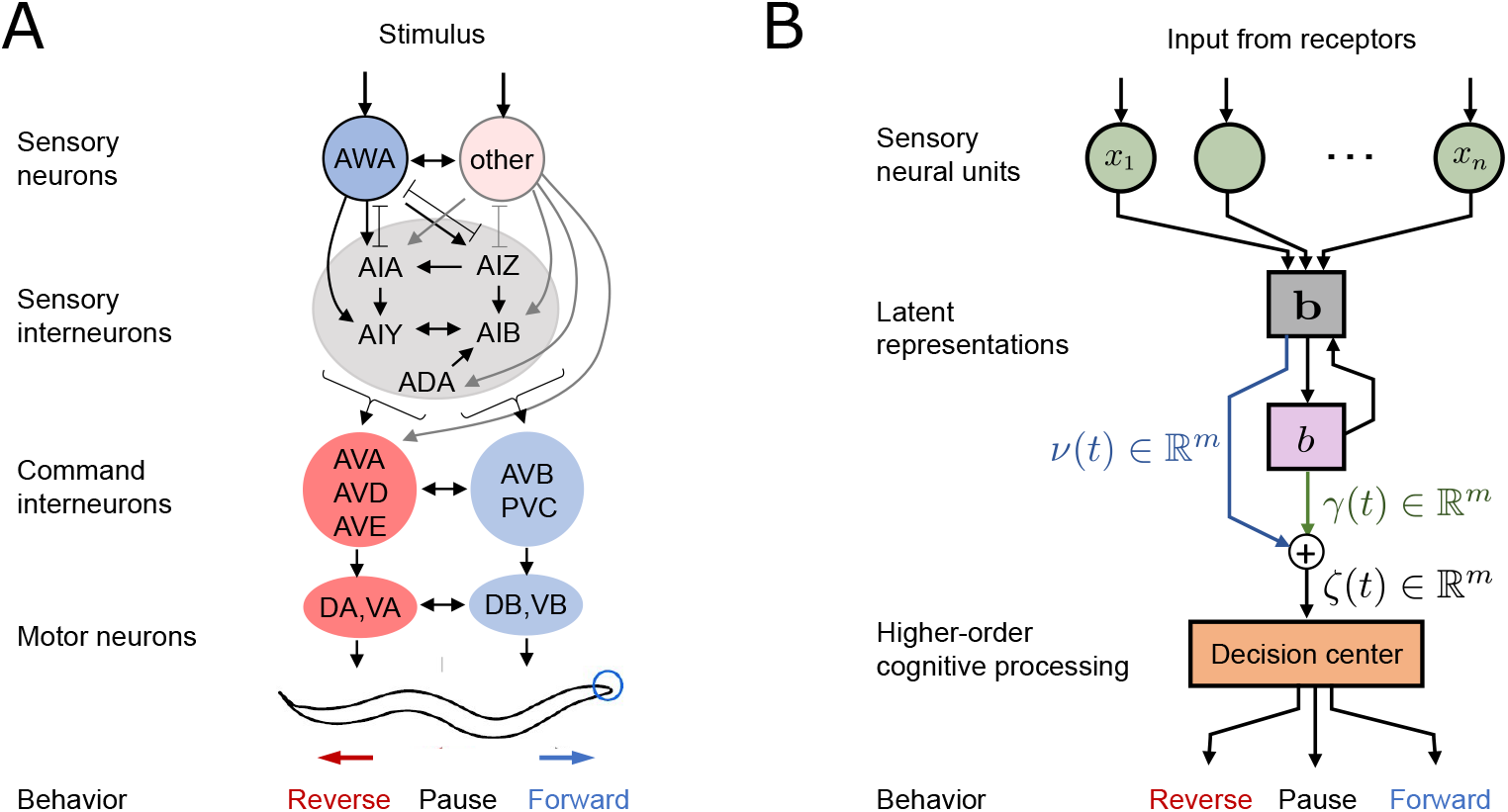
Information flow from stimulus to behavior. **A**. Neural block diagram of sensory circuit from afferent sensory to effector neurons. **B**. Schematic depicting signal flow in the computational model. Note that while intermediate variables are mathematically abstracted, they can be interpreted as activity of interneurons.

The computational model is built by drawing on a classical problem in control systems theory, the Finite Horizon Linear Quadratic Regulator (LQR) (e.g., ***Anderson and Moore (2007***)). The central theoretical premise is the assumption that sensory neural activity drives the creation of an actionable latent representation, formed downstream. The process of creating this representation is governed by a linear dynamical systems model, whose dynamics are comparable to classical drift-diffusion type decoders found in detection and decision models ***Mallik et al. (2020a***); ***Mormann et al. (2010***); ***Veliz-Cuba et al. (2016***). By postulating certain priorities regarding the speed and fidelity by which representations are constructed, we can thus synthesize and predict the driving neural activity and associated neural dynamics. Specifically, we use the following mathematical problem, which minimizes cost function 𝕁(**x**) penalizing three factors: (i) inaccuracy of the fast latent representation (via **Q**); (ii) energy expenditure during neural activation (via **S**); and (iii) rapid fluctuations in neural response (via **R**):

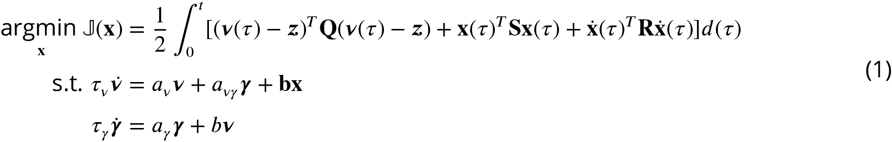

Here, ***z*** is a nominal ‘target’ sensory representation that corresponds to a specific cue from the periphery, while ***v*** is the actual fast-evolving latent representation formed from neural activity that is responsible for mediating immediate sensory detection and response (Figure 3A). Further, ***γ*** is a slow evolving latent representation representing short-term memory, while **x** represents activity of the sensory neurons. The parameters *τ*_*v*_, *τ*_*y*_ are the timescales of the immediate and slow latent representations. Coefficients *a*_*v*_ *<* 0, *a*_*vγ*_ ∈ℝ, *a*_*γ*_ *<* 0 represent the dynamics of the decoder that map neural activity **x**) into latent representations. Matrix **b** weights the contribution of neural unit activity and projects it onto the latent space dynamics. Finally, *b* scales the contribution of the immediate latent representation onto the dynamics of residual memory. A detailed technical account of how we conceptualized this optimization problem and chose the parameters is provided in the Methods section. Briefly, the cost function 𝕁(**x**) is optimized when neural activity **x** efficiently, smoothly and quickly creates actionable latent representations. The optimal dynamics for **x** is obtained by solving (1), yielding:

**Figure 3.**
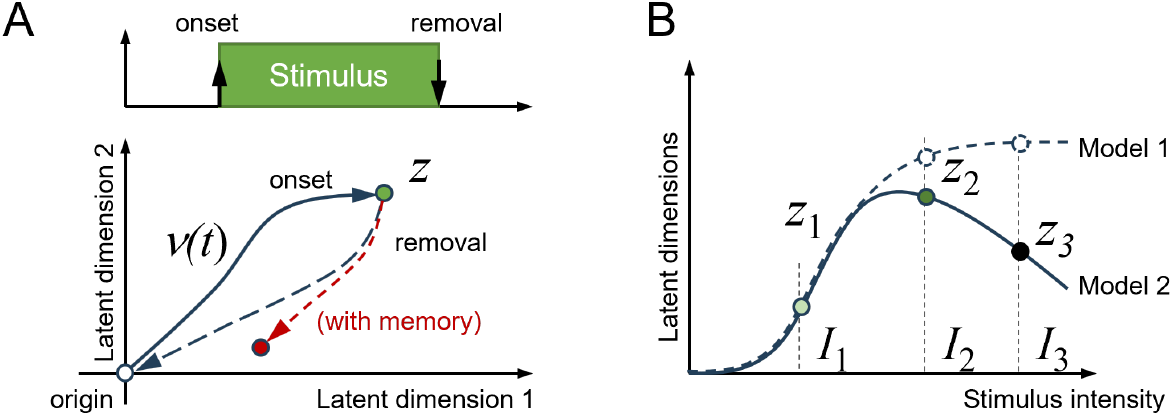
Latent representations of stimulus onset and removal. **A**. In presence of stimulus, the latent trajectory evolves starting from origin or the neutral regime to track ***z***. When the stimulus is withdrawn, the latent trajectory promptly returns to the neutral regime, either at the origin or at a new latent state if residual memory is established. **B**. The odor stimulus is characterized by its identity and intensity. In Model 1, latent representations increase monotonically with intensity, whereas in Model 2, latent representations peak at intermediate intensities.

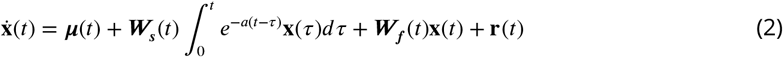

Here, the first term on the right hand side ***µ***(*t*) represents feedback from the memory representation onto sensory neural dynamics (see in Methods and Appendix 1 the details for derivation). The second and third terms represent slow and fast processing of neural response respectively. The final term **r**(*t*) represents the afferent inputs that the sensory networks receive from upstream receptors. The matrices ***W***_*s*_(*t*) and ***W***_*f*_ (*t*) provide the weights of the connections. Note that in the limit of **x** describing a single neuron, then these matrices reduce to scalars. The mathematical details of how we derive (2) follows from our previous work ***Mallik et al. (2020a***) and the key steps are outlined in Appendix 1. We note here that **x** is a dimensionless variable within this mathematical problem, and in order to compare the results of model simulation with experimental fluorescence imaging recordings, we add post-processing steps that mirror the experimental process (see Figure 8 below).

<H23Model part II: Stimulus intensity and preferred latent representation

In our model, stimulus intensity is encoded in the nominal target representation ***z***. We consider two models of intensity embedding (see Figure 3B):

*Model 1: Monotonic scaling*, where the target latent representation monotonically moves away from the neutral regime of the latent space (i.e., *v* = 0) and saturates away from the neutral regime beyond a certain intensity, and

*Model 2: Preferred intensity*, where the target latent representation initially moves away from the origin and reverses course beyond a certain intensity, resulting in a ‘preferred’ maximal intensity.

Organisms inherently exhibit preference to certain stimulus identities and intensities, and *C*.*elegans* prefer intermediate conditions (such as salt, temperature, and oxygen levels) that were present during cultivation or recent beneficial feeding ***Bargmann (2006b***); ***Matsumoto et al. (2024***); ***Chang et al. (2006***); ***Almoril-Porras et al. (2025***); ***Kimata et al. (2012***). Highly attractive stimuli often become aversive at high concentrations ***Lagoy (2018***); ***Zhang et al. (2024***). Hence, we used Model 2 of intensity encoding to better match these observations.

#### Model part III: Generating behavior through a Bayesian decoder

The latent representation constructed in our framework constitutes a fluid encoding of the periphery, fluctuating in response to sensory cues, and actionable when routed to higher brain regions for behavior generation. As such, we funnel the latent representations into a Bayesian decoder to motor outputs corresponding to probability of forward, pause, and reversal behaviors in *C. elegans* (see **Methods** for details). The Bayesian decoder in this work is parametric and requires specification of nominal attractive representations resulting in forward motor behavior. We used the *C. elegans* appetitive food odorant diacetyl as a reference for an attractive stimulus.

Using the optimization framework, we parameterized the neural response model and the Bayesian behavior decoder to match experimental responses to one stimulus concentration (4.4 *µ*M diacetyl) and timing pattern (20 s stimulus every 60 s). Simulated neural and behavioral responses for this fixed intensity captured neural adaptation and behavioral invariance (Figure 4). As the stimulus repeats, the organism through its neural activity should generate latent representations that maintain the same nominal target. When sensory memory is actively engaged in sequential stimulus detection, neural activity decreases exponentially with repeated encounters (Figure 4A), which matches experimental adaptation used for optimization (Figure 4B). The fast latent representation *v* closely follows the stimulus pattern, with dimensions 1 and 2 representing the low- and highpass filtered input signal (time constant *r* 1 sec), respectively (Figure 4C). On the other hand, the involvement of memory in the generation of motor output results in behavioral invariance (see Figure 4D,F,G). After rising during several initial stimulus pulses, a steady level of forward behavior is seen across subsequent trials indicating invariant behavior without habituation, despite neural adaptation. These model predictions match experimental results for behavior, which were used to parameterize the Bayesian decoder (Figure 4E).

**Figure 4.**
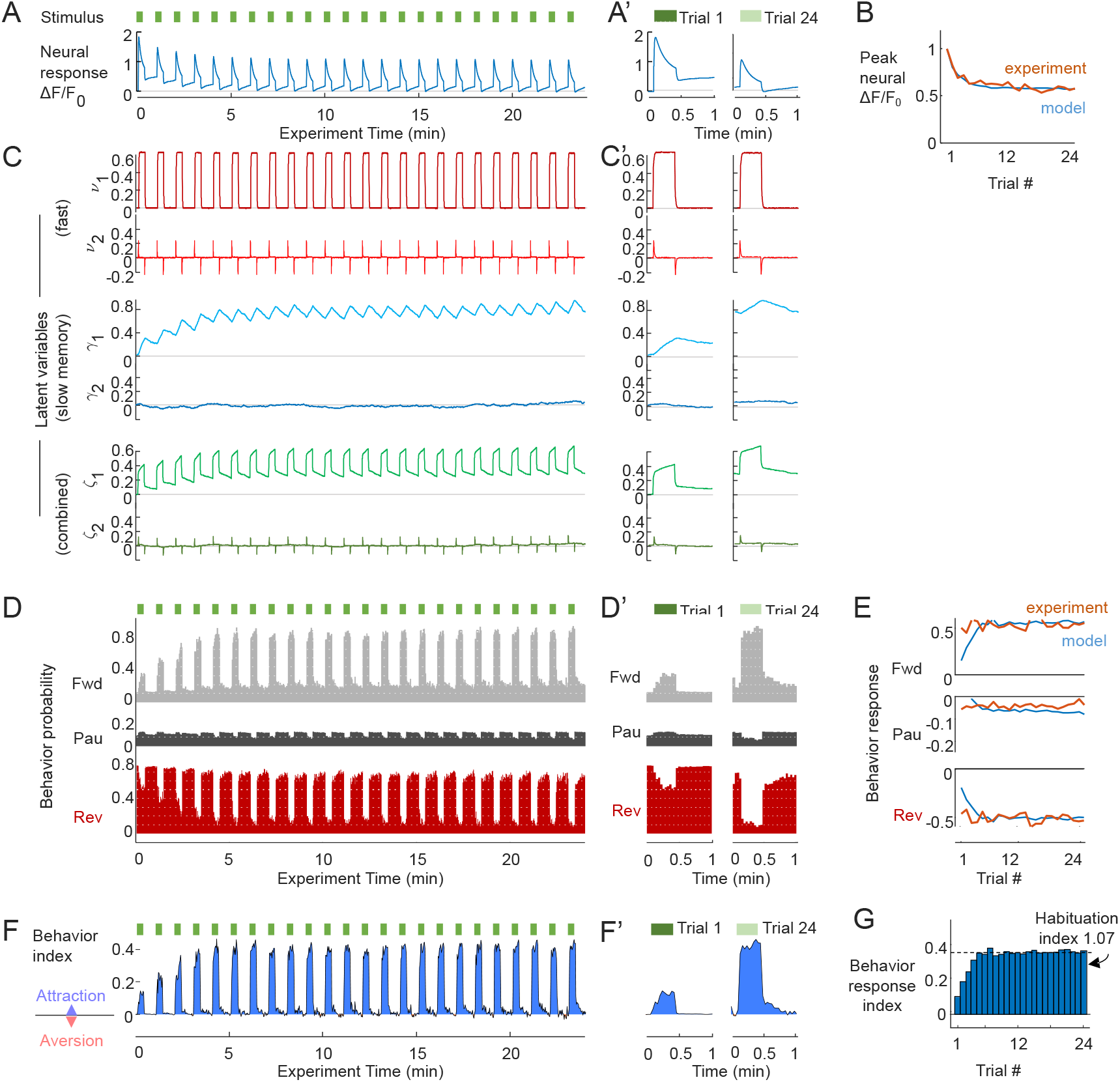
**A**. Simulated neural calcium responses to 20s stimulation pulses every 60s (green bars). Trials 1 and 24 are magnified in panel A’. **B**. Peak neural calcium over 24 repeated trials, normalized to the initial peak,comparing simulations (blue) with experimental measurements of AWA responses to 4.4 *µM* diacetyl odor (orange). Relative peak amplitude adapts exponentially and reaches a steady asymptote after about 5 repeated encounters in both model and experiment. **C**. Corresponding latent variables: fast ***v***(*t*), slow memory ***γ*** (*t*), and combination *ζ*(*t*) = (***v*** + ***γ***)/2. Trials 1 and 24 are magnified in panel C’. **D**. Simulated behavioral probability for Forward (Fwd), Pause (Pau) and Reverse (Rev) states show consistent increase in forward locomotion during stimulation (green bars), and decrease in pause and reverse state probability. Trials 1 and 24 are magnified in panel D’. **E**. Behavioral response probability for forward, pause, and reverse states across 24 repeated stimuli, defined as mean state probability change during stimulation vs. non-stimulation. Behavioral responses for 20s pulses remain invariant in both model and experiment. **F**. The Behavior index indicates instantaneous attractive behavior by positive values and aversive responses by negative values. For 20 s stimuli, response predictions are consistently attractive. Trials 1 and 24 are magnified in F’. **G**. The Behavior response index quantifies the attractiveness of each trial’s response, defined as mean forward Behavior index during stimulation vs. non-stimulation. The Habituation index quantifies degree of habituation as the response index at the ending trials versus the initial trials. A Habituation index of 0 indicates complete habituation (no response) whereas a value of 1 indicates behavioral invariance.

### Stimulus timing inluences neural adaptation: prediction & validation

To further validate these model interpretations, we sought to make predictions regarding the neural and behavioral responses of animals to previously untested stimulus scenarios.

We first examined the model with regard to different stimulation timing patterns, without changing the model parameters. We sought to validate the model’s predictions of the rate and asymptotic level of neural adaptation. Periodic stimulus pulses are defined by pulse duration (PD), intertrial interval (ITI, between pulse initiations), and their ratio, the duty cycle (DC, or time fraction of stimulus presence):

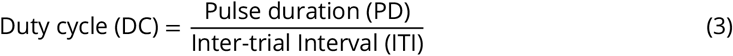

Beginning with the central stimulation timing (PD = 20s, ITI = 60s, DC = 33%), we simulated eight additional stimulation patterns both increasing and decreasing PD, ITI, and DC by 2.5-fold (see Figure 5A). This scheme enables comparison of neural response trends across each variable. We emphasize that for the additional timing patterns, no further model optimization was performed.

**Figure 5.**
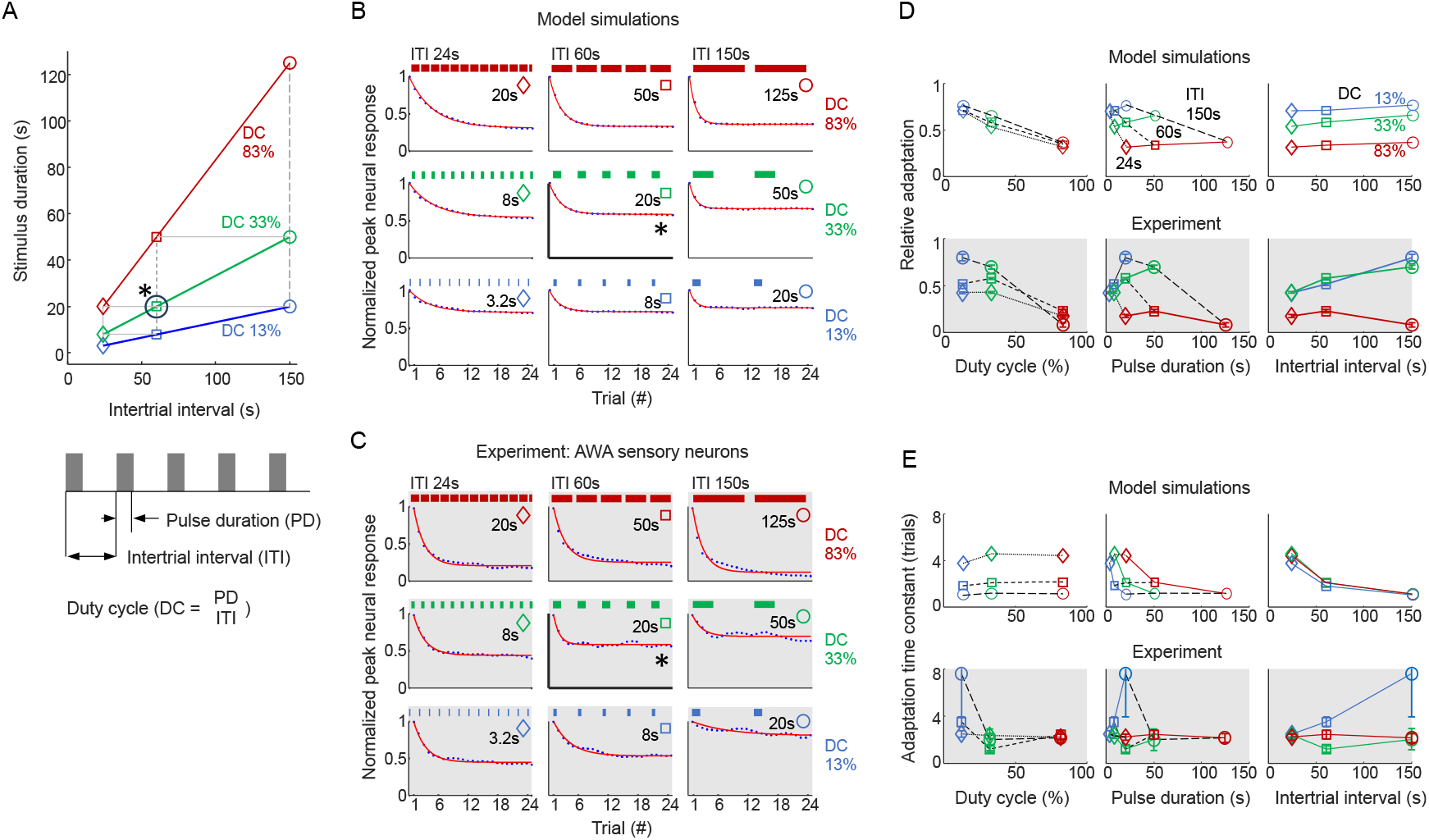
Adaptation rate and level depend on stimulation timing. **A**. Stimulation paradigm allowing comparison across duty cycle, pulse duration and inter-trial interval. **B**. Model simulation of relative peak activity for a representative animal. Curves represent exponential fits as described in Statistical Methods. A representation of the stimulus timing is shown above each graph. **C**. Experimental measurements of the *C. elegans* AWA neuron peak response per trial presenting 4.4 *µM* diacetyl pulses. **D**. Relative adaptation level (trial 24 vs. naive trial 1 neural response) depicted for model simulation (above) and experiment (below), indicating good model prediction. **E**. Adaptation rate (time constant *τ* in units of trial number), depicted for model simulation of a representative animal from B (above) and experiment (below), is less well predicted by the model. Error bars shown in D,E are the 95% confidence interval of each value, n = 18-23 animals.

We observed the following model predictions (Figure 5B) and note which match experimental results (Figure 5C):

- The model predicts that peak neural responses decay exponentially following repeated stimulus exposures, to a steady asymptotic adaptation level dependent primarily on the *duty cycle* (DC) of the stimulus. Higher DC leads to greater neural adaptation, and experimental data validate this prediction (Figure 5D).
- The model predicts that pulse duration (PD) and inter-trial interval (ITI) do not directly contribute to adaptation level in the model, except as they contribute to DC. Experimental results are similar, although adaptation levels are higher for longer ITI at moderate DC (Figure 5D).
- The model predicts that ITI should contribute to adaptation rate (*τ*), with higher ITIs adapting faster. Experiments do *not* match these predictions.

Thus, with model optimization to a single stimulation pattern, simulations correctly predict the level of adaptation for different duty cycles, pulse durations, and inter-trial intervals, but adaptation timing varies less in *C. elegans* sensory neurons than in model predictions.

### Behavioral invariance, habituation, and inversion: prediction & validation

We next interrogated our model with respect to stimulus intensity, another key factor that influences dynamically evolving latent information.

We simulated habituation across 24 repeated trials across a range of stimulus intensity and duty cycle for a fixed ITI, calculating for each condition a Habituation Index (Figure 6A). Three predictions were apparent regarding the neural-behavioral mapping over repeated stimuli:

**Figure 6.**
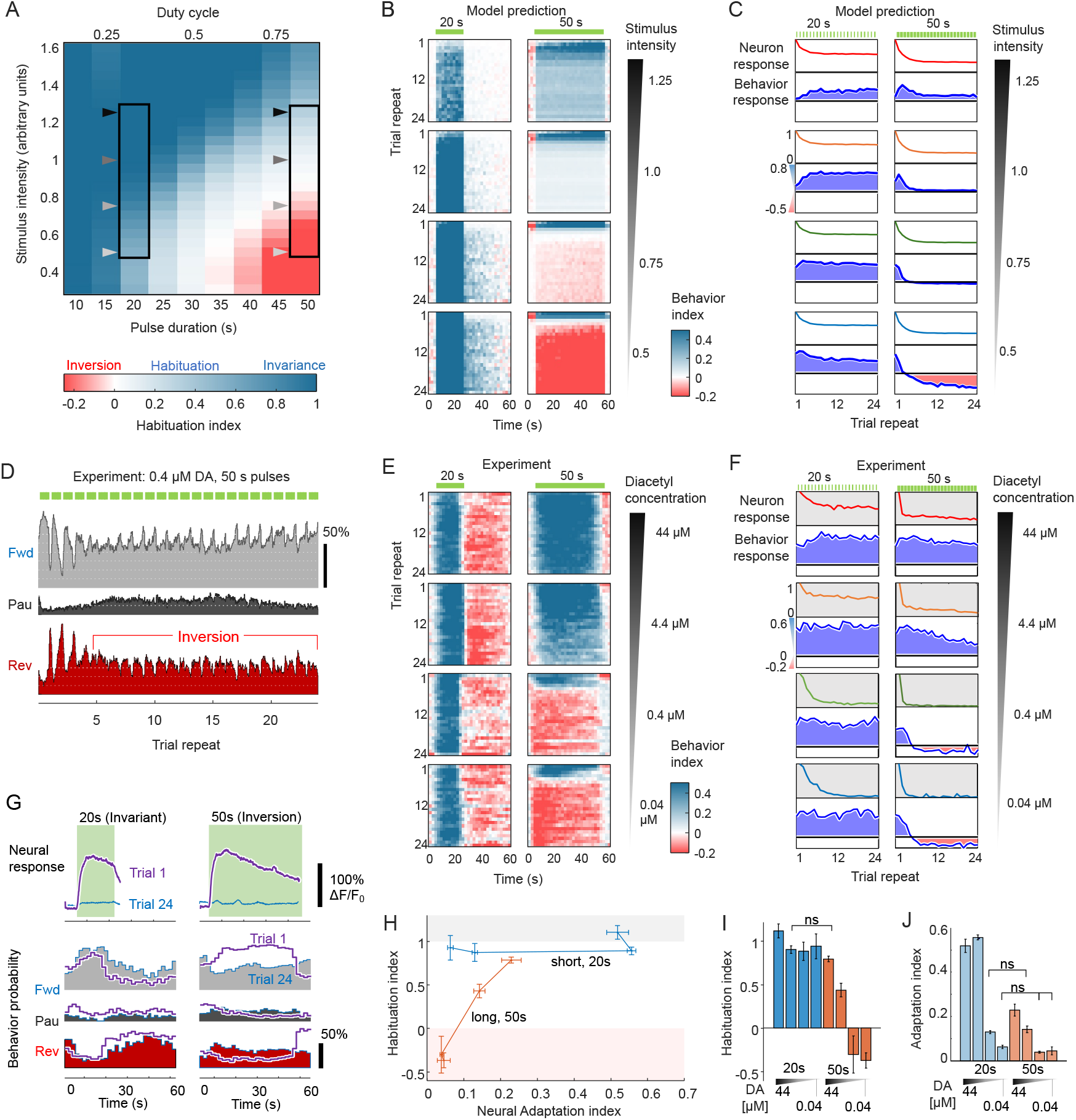
Stimulus intensity and duration determine habituation. **A**. Model predictions of habituation over 24 repeated stimuli for varying stimulus intensity and duration (60 s ISI). Heatmap shows Habituation Index (HI), defined as the normalized behavior Response Index of Trial 24 versus Trial 1. Behavioral invariance (dark blue, HI ≈1) occurs high stimulus intensity, whereas habituation (light blue, 0≤HI<1) is predicted at lower stimulus intensity and longer durations. Behavioral inversion (red, HI<1) is predicted for very long, low intensity stimulation. Arrowheads indicate model conditions depicted in panels B and C. **B**. Model predictions of Behavior Index during each 60s trial for four selected intensities and two durations, 20s and 50s. Whereas 20s stimuli elicit attractive behavior responses (blue) at all concentrations, 50s responses are initially attractive but habituate or invert after a few trials at lower intensities. **C**. Model predictions of peak neural response and Behavior response index for the intensity and duration conditions in panel B. For short 20s stimuli, behavior is predicted to be invariant irrespective of stimulus intensity, despite 2-fold neural adaptation. For long 50s stimuli, behavioral predictions show slight habituation at higher intensities and inversion at lower intensities c < 0.75. **D**. Experimental validation of behavioral inversion. Behavior probability in response to long 0.4 *µ*M diacetyl pulses (50s duration) indicates an attractive response for the first four stimuli (elevated forward probability during stimulation, green bars) followed by aversive responses. Plots indicate average instantaneous state probability, n = 107 animals. **E**. Experimental measurement of attractive Behavior Index for four different intensities, 0.04 - 44 *µ*M, at pulse durations 20s and 50s. For 20s pulse duration, behavior remains invariant across trials, with attractive behavior elicited during stimulation (blue) and suppressed upon stimulus removal (red). Similar results are observed for 50 s pulses at high concentration, whereas at low intensity, behavioral inversion occurs after several stimuli. All of these results match model predictions (panel C). Data are averaged across 28 - 107 animals. **F**. Experimental peak neural and behavioral response index for conditions identified in E. **G**. Mean neural and behavioral responses for Trials 1 and 24 for short (20 s) and long (50 s) pulses of 0.04 *µ*M diacetyl stimulus. Neural adaptation is equally strong for both stimulus durations, but habituation differs between short stimuli (behavioral invariance) and long stimuli (inversion). n = 28,23 animals (neural 20s, 50s) and n = 30,90 animals (behavior 20s, 50s). **H**. Plot of Habituation index versus Neural adaptation index for long and short pulse experiments at four concentrations, increasing in the direction of arrows. Error bars indicate standard deviation of indices. Habituation index **I** and Adaptation index **J** for short (blue) and long (red) pulses at the indicated range of stimulus concentration. ns, not significant (p>0.05).

- First, no or minimal habituation (behavioral invariance) is predicted for shorter duty cycles (DC 33%) at all stimulus intensities (see Figure 6A). Under these conditions, equal increase in forward probability is predicted across repeated trials, indicating an invariant attractive response.
- Second, at greater duty cycles 33%, behavior begins to habituate to repeated encounters of longer stimulus durations, albeit at a much slower rate than that observed in neural adaptation (see Figure 6). This occurs in a dose-dependent manner.
- Third, and perhaps more intriguingly, simulations predict at high duty cycle (DC > 66%) the emergence of a paradoxical phenomenon. Under these conditions, early responses reflect the correct stimulus representation, i.e. appetitive stimuli elicit increased forward behaviors. These forward responses habituate during the next few trials, then *invert* to a weakly aversive response (i.e., more reversals, Figure 6B,C). We call this switch in response valence **behavioral inversion**.

To explore these predictions of behavioral invariance, habituation, and inversion, we experimentally stimulated wild-type *C. elegans* 24 times (once per minute) with short (20 s duration, DC = 33%) and long (50 s duration, DC = 83%) pulses of diacetyl odor at four concentrations spanning three orders of magnitude from 44 nM to 44 *µ*M, representing a range of stimulus concentration responses from weak but detectable to very strong ***Lagoy (2018***). All three predictions were observed:

- Experimental behavioral responses were indeed invariant for short 20 s pulses at all concentrations (Prediction 1; Figure 6E).
- Long 50 s pulses also elicited invariant behavior at the highest stimulus concentration, whereas the lower 4 *µ*M concentration elicited steadily habituating responses over time (Prediction 2; Figure 6E,F).
- Long 50 s pulses of the lowest concentrations demonstrated initially strong behavioral responses that habituated over three to four trials, then inverted for the remaining trials (Prediction 3; Figure 6D-F). Hence, this paradoxical “behavioral inversion” model prediction in fact occurs *in vivo*.

At low stimulus intensity (0.4 *µ*M), short 20-s and long 50-s pulses elicit neural responses are indistinguishable in peak levels and dynamics during stimulation at both the first and last repeated trials (Figure 6G); indeed, both have similar neural Adaptation indices (0.063 ± 0.008 vs. 0.045 ± 0.018, respectively, P>0.16). Yet, differences in behavioral response are apparent immediately at stimulus onset, with consistently attractive responses for short stimuli, and inverting responses for long stimuli, beginning within seconds of stimulation. Thus, neural information alone does not predict behavioral outcome. Under conditions leading to strong adaptation (Adaptation index < 0.25), habituation can span the full range from invariance to inversion (Figure 6H-J).

## Discussion

### Short-term latent memory traces account for neural to behavioral transformations

In our study, we investigated the sequential mapping from a peripheral sensory stimulus to a sensory neural response, culminating in a behavioral output. Our results demonstrate that this latter transformation is not injective. That is, it is likely not possible to simply map a level of sensory neural response to particular behavioral probabilities (like, for instance, lower sensorimotor-to-behavior coupling ***Atanas et al. (2023***)). This conclusion is immediately apparent in light of our observation of behavioral invariance, wherein animals are seemingly able to maintain very stable behavioral responses despite waning neural activity due to adaptation.

We posit an explanation for these observations based on the theory that organisms are constructing a latent stimulus representation that persists and evolves on a slower time scale than that of the immediate sensory neural response. The presence of such a representation, a form of shortterm sensory memory, predicts many features of neural adaptation to repetitive stimuli, which we validate by varying the stimulus dynamics and characterizing neural and behavioral outputs. Our computational model predicted the levels of experimental neural adaptation well, without changing its parameters, with greater adaptation for a longer duty cycle and shorter trial intervals. More intriguingly, paradoxical regimes were predicted in which behavioral responses can invert over repetitions, relative to the nominal peripheral stimulus that is present. Taken together, our integration of theory and experiment affirms a key role for dynamical processes beyond overt sensory neural activity in the creation of contextual, history-dependent behavior.

### Multiple time-scales and behavioral inversion

#### Theoretical premise

The aforementioned behavioral inversion phenomenon is especially interesting because it is emergent in the theoretical model under specific stimulus conditions (i.e., low intensity, long stimulus durations). It is useful to review the flow of signals through our model and especially the interpretation of the variable *ζ*, the aggregated signal used by the Bayesian behavioral decoder. This variable is a linear combination of latent representations evolving along fast (i.e., ***v***) and slow (i.e., memory, ***γ***) time-scales. As depicted in Figure 4C, the dimensions of *ζ* correspond to behaviorally salient features. In our study, we observe that there is a rapid evolution of *ζ* from the stimulus that occurs over the first approximately two seconds, quickly rising along dimension 1, and rising then falling along dimension 2 (Fig 7A). During stimulation, *ζ*gradually increases, drifting until the moment of stimulus removal. The return path follows similar kinetics, inverted along dimension However, the latent representation does not return to the origin starting point, but rather shifts by an amount similar to the drift during stimulus presentation. This indicates a hysteretic buildup of slow sensory memory (Figure 3A). If we examine the stimulus-on kinetics of the first, third and twenty-fourth trials, we observe that low concentrations and large duty cycles push the latent variable curves positively along the *ζ*_1_ axis (Figure 7B). Behavior valence is largely determined by *ζ*_1_ (Figure 7C):

**Figure 7.**
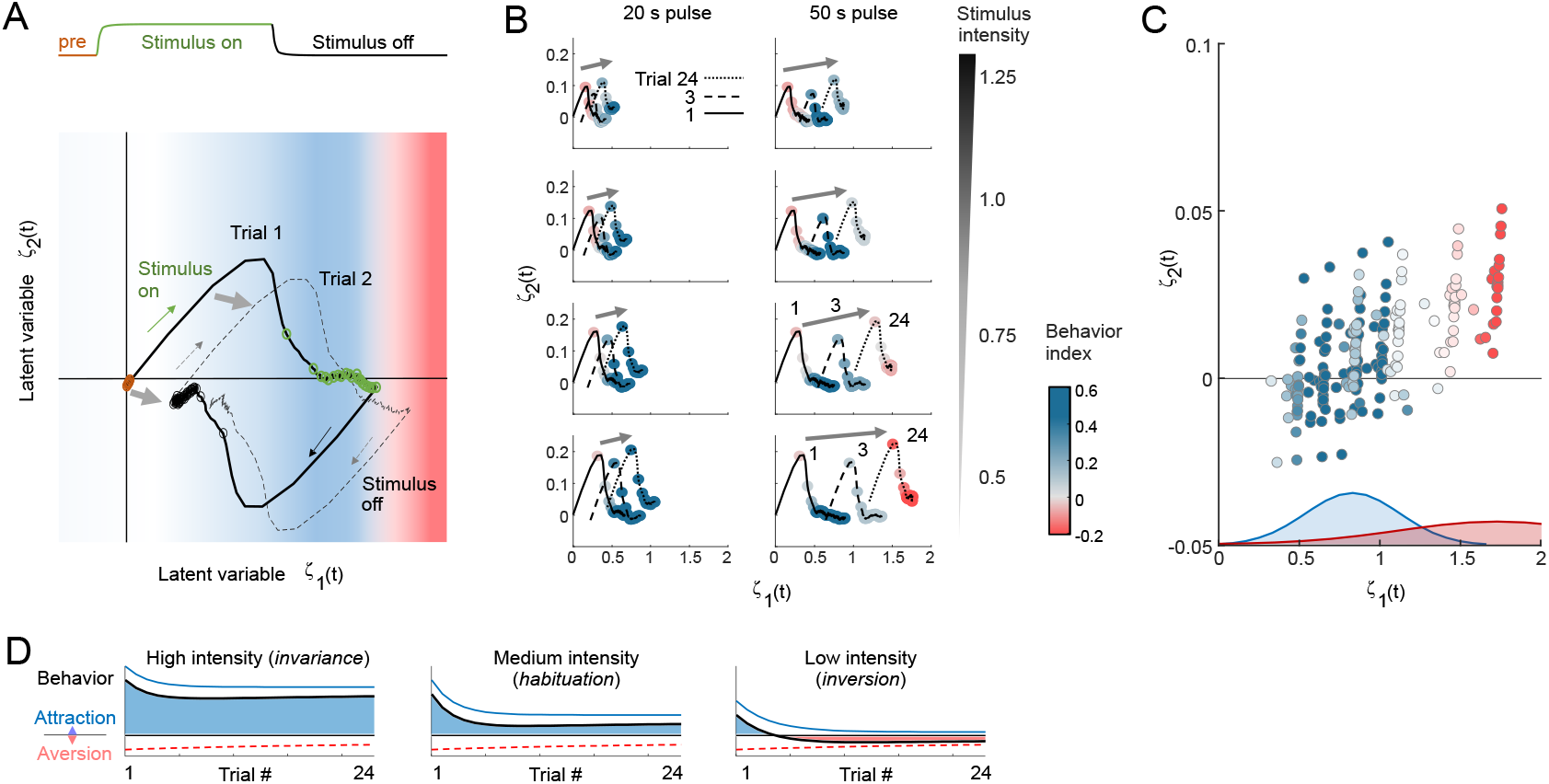
**A**. Schematic depicting the two-dimensional latent space and a weighted combination of latent representations ***v*** and ***r*** evolving in presence of stimulus across two trials. Latent dimension 1 represents the primary stimulus representation whereas dimension 2 represents ambiguity arising from the sudden onset or offset of stimulus. Notice that the latent variable curves shift each trial, increasing primarily in dimension 1 and to a lesser extent in dimension 2 (gray arrows). Curve is plotted for intensity c = 1.0, short 20 s pulse. Open circles represent 1 s time intervals (prestimulus = orange, stimulus on = green, stimulus off = black). **B**. Latent representations are plotted for short (20 s) and long (50 s) stimulus pulses at low to high stimulus intensity. Trials 1, 3, and 24 are shown for stimulus time points only. Colored timepoints represent the attractive Behavior index. At low concentrations and long pulses, curves shift leftward leading to a region of behavioral inversion. **C**. Latent representations at the end of stimulation for all trials shows behavioral inversion occurring for *ζ*_1_ > 1.5. Here, *ζ*_2_ plays little apparent role in behavior response. Valence thresholds arise through parameterization of the Bayesian decoder, which specifies mean and variance of Gaussian shapes for appetitive behavior (forward, blue shading) and aversive behaviors (pause and reverse, red shading). **D**. Hypothetical summation of attractive and aversive signals evoked by the same stimulus and adapting to repeated trials at different, intensity-dependent, rates and levels. Here, invariance (high intensity), habituation (medium intensity), and inversion (low intensity) depend on the relative strength of attractive and aversive signals.

**Figure 8.**
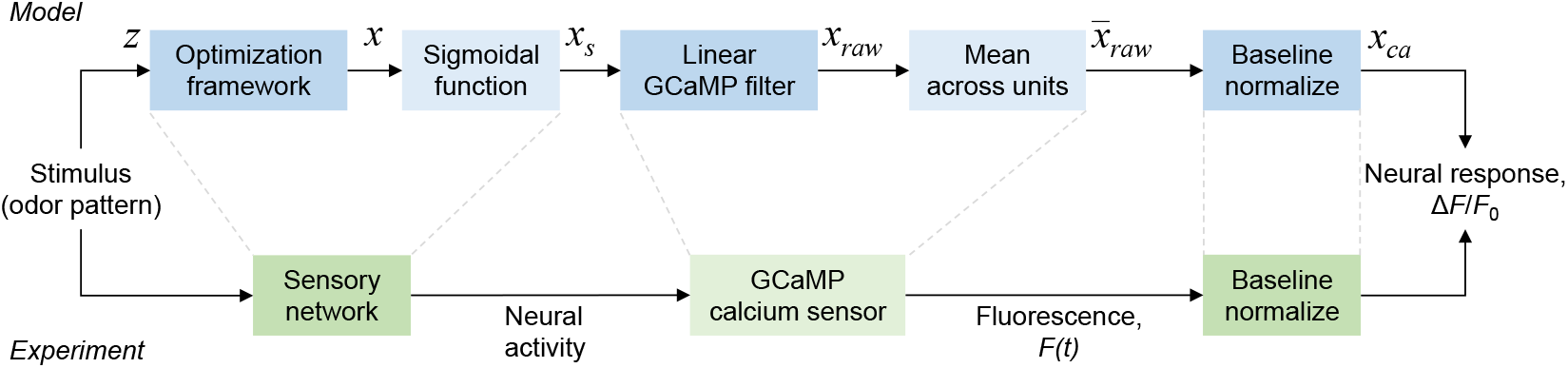
Post-processing pipeline

- between 0 and 1, responses to repeated stimulation are invariant;
- from 1 to 1.5, responses habituate; and
- above 1.5, behavioral inversion occurs.

The behavioral transition point of several positive responses before inversion occurs can be explained by the slow accumulation of *ζ*_1_ to reach the threshold values for inversion only after 3 - 4 trials (see Figure 6D and 7B). Conceptually, these threshold values arise from the overlay of multiple Gaussian distributions. Interpreted in its simplest form, the summation of an attractive distribution and an aversive distribution with a wider variance would produce the observed thresholds (Figure 7C).

#### In our model, behavioral inversion emerges from the interplay of the following factors

1. The behavior model parameters, which specify the likelihood of attractive or aversive behavior responses given latent representation of the stimulus and were fit to experimental data (Figure 4B,E). The mean and variance of Gaussian distributions for attractive and aversive behavior probability sum linearly, leading to the valence threshold where inversion occurs when *ζ*_1_ > 1.5 (Figure 7C).
2. The accumulation of short-term memory in the latent representation, which increases more during longer stimulation durations.
3. The scaling of target latent preference of the animal, encoded in the preferred stimulus intensity model *z*_*c*_ (see Figure 3B and equation 9). Intensity model parameters determine the concentrations at which inversion occurs (Appendix 1 Figure 1), and were selected to match experimental data. Here, we found the correct prediction occurs when *z*_*c*_ *decreases* with concentration (Appendix 1 Figure 1B and Figure 7B).
4. The logarithmic mapping of experiment stimulus concentration to model concentration, representing fold-change similarity in behavior ***Larsch et al. (2015***). Here, we use the relation 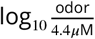 such that *c* = 1.0 reflects the base optimization stimulus and *c* increases with odor concentration.

Experimentally, inversion was observed only for very low odor concentrations. Therefore, the preferred stimulus model required *z*_*c*_ to be greatest at these lower concentrations, and hence latent representation *z* does not strictly scale with scalar stimulus intensity. Instead, it may represent the summation of several stimulus-evoked signals of opposing valence, each having its own adaptation level and rate. For example, we hypothesize that appetitive signals scale with attractive odor intensity and adapt quickly, whereas aversive signals are weaker and adapt more slowly. Under these conditions, the summation of signals yields invariance at high odor concentrations and inversion after a few trials at low concentration (Figure 7D).

#### Mechanistic interpretations

Where in the nervous system are latent representations encoded, if not within sensory neural dynamics? While our results do not rule out sensory dynamics as drivers of habituation (for example, sensory activity is prolonged during longer stimuli, see Figure 6G), they suggest latent variables are encoded elsewhere. A network of neurons in the chemosensory circuit of *C. elegans* carries out complex neural to behavioral transformations, including sensory interneurons, command interneurons, and motorneurons ***Bargmann (2006a***); ***Gray et al. (2005***); ***White et al. (1986***), and other neurons sensitive to the stimulus could provide additional inputs ***Lagoy (2018***); ***Dobosiewicz et al. (2019***). For example, in *C. elegans* an AWC-detected odorant perceived as attractive in isolation can become aversive when presented alongside other attractants ***Khan et al. (2022***), and the multimodal ASE neuron can trigger opposite chemotactic responses by engaging different interneuron circuits ***Xue et al. (2025***). In mammals, the nucleus accumbens has shown similar capability to flexibly assign stimulus valence, although how this occurs remains under active investigation ***Vieitas-Gaspar et al. (2025***). Co-activation of multiple neurons, perhaps with opposite valence, could lead to the behavioral inversion phenotype, yet this scenario is unlikely as inversion occurs at low stimulus concentration, and the AWA neurons we studied are the most sensitive to the diacetyl stimulus ***Sengupta et al. (1996***). In the future, whole brain imaging studies can help elucidate other neural units that contribute to the higher dimensionality of latent representations, especially with the recent introduction of high-fidelity transgenic constructs for the quantification of responses and their associated neuron identities ***Yemini et al. (2021***); ***Atanas et al. (2023***).

An intriguing alternative is the role of neuromodulators in these representations ***Flavell et al.(2022***), such as neuropeptides that regulate neural adaptation ***Chalasani et al. (2010***) and behavioral habituation to other stimuli ***Ardiel et al. (2017***). Pre-synaptic neural activation can co-release local, rapid, excitatory neurotransmitters and diffuse, slower, inhibitory neuropeptide signals ***Méndez-***

***Couz et al. (2021***). Hence, aversive behavioral responses to attractive stimuli could hypothetically arise when an inhibitory neuropeptide signal exceeds the excitatory neurotransmitter signal (as in Figure 7D). Future experiments will address this hypothesis in *C. elegans* using neuropeptide-deficient animals.

### Limitations and Features not Explained

#### Within-trial model response dynamics

The focus of our modeling was on the adaptation of the magnitude of neural and behavioral responses to repeated stimulation. Although the model predicts neural response dynamics well, it is less accurate during stimulus removal between individual stimulus trials. Whereas experimental recordings decay exponentially to baseline levels within <30 s, model output does not reach baseline in early trials and increases slightly before each subsequent stimulation (compare Figures 1B and 4A). We note that optimization can produce better fit to the neural dynamics of individual trials, but this comes at the expense of accuracy of peak response levels during adaptation. This suggests room for improvement with additional neural model parameters. Similarly, optimization of the behavioral output for the Bayesian decoder was performed to capture the overall trends of stimulus onset and offset behavior for the whole experiment, rather than exact temporal dynamics. Additional parameters and optimization may more closely match within-trial behavioral dynamics.

#### Responses to non-uniform stimulation and memory decay

Here, we simulated and measured responses to stimulus pulses with uniform timing and intensity. We did not validate how well the model predicts the recovery from adaptation and habituation when stimulation is removed. Further, we have not tested adaptation and habituation rates to more complex, naturalistic stimulus environments, such as sensory information that ebbs and flows over time during chemotaxis behavior toward a food source. Methods are available to measure neural and behavioral responses to varied stimulus intensity levels and timing within a continuous trial ***White et al. (2023***), and these results will be useful to further refine the model presented here.

### Circuit-level implementation of the model

Our results constitute a hypothesis for the dynamical mechanisms and factors that mediate the observed neural and behavioral relationships. However, our theoretical work does not assert specific neuronal pathways that might instantiate these computations. One might imagine that hierarchical processing through neurons and synapses might also play a role in representing sensory stimuli across timescales. Clearly, the next steps for this work involve a more circuit-oriented investigation of these mechanisms, including examination of a larger assortment of other sensory neurons, such as those that encode aversive stimuli).

## Methods and Materials

### Computational Methods

#### Latent representations created by sensory activity

Previous theoretical and experimental studies ***Sadtler et al. (2014***); ***Cunningham and Byron (2014***) have suggested that information accrued by sensory systems can be represented in a compact, lowdimensional format. In this premise, we introduce the concept of a latent space where stimulus specific representations are created dynamically. We posit that activity of sensory neurons drive manifestations in this latent space. Intuitively, the latent space is akin to a high-level feature space, with each dimension representing some crude, stimulus specific feature such as stimulus identity, valence, intensity, novelty, or urgency.

In our model, we represent by **x**(*t*) ∈ ℝ^*n*^ the instantaneous activity of sensory neural units (with *n* being the number of units considered). The activity of sensory neural units aid in the creation of latent representation along two different timescales - (a) representation associated with immediate detection: ***ν***(*t*) ∈ ℝ. (b) representation associated with residual memory: ***γ*** (*t*) ∈ ℝ. Here, is the dimensionality of the latent space and *m*≤ *n*. More specifically, the temporal evolution of latent representations can be expressed mathematically as:

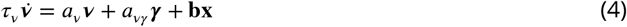

and,

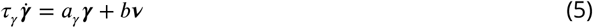

Here, *τ*_*v*_, *τ*_*y*_ are the timescales of latent representations for immediate detection and residual memory respectively. *a*_*v*_ *<* 0, *a*_*vy*_ ∈ ℝ, *a*_*y*_ *<* 0 are the coefficients representing internal decoder dynamics. **b** weights the contribution of neural unit activity and projects it onto the latent space dynamics. Finally, *b* scales the contribution of the immediate latent representation in the dynamics of residual memory. The parameters of this model are inferred by minimizing RMSE between simulated response and experimental recordings (see in Appendix 1).

Note that this is a generalized mathematical framework for stimulus information encoding; hence, *n* in our setup is an arbitrary positive integer. Specifically for the AWA chemosensory neurons, we choose *n* = 2 to represent either the left-right neuron pair (which can respond differently, but usually respond equally) or the combination of direct neural activation and feedback signals from other neurons (such as aversive neuron ASH, which can suppress AWA activity ***Lagoy (2018***)).

### Box 1. Theoretical and experimental worklow

In this work, we have explored neural and behavioral adaptation through theoretical and experimental methods. More specifically, experimentation and computational models are complementary components in our study. We also introduce a prediction validation scheme through which we characterize the dependence of neural and behavioral activity on external stimulus parameters. In Figure 1, we have outlined the work-flow for this study and how comparisons were made between experimentally observed quantities and simulated variables.

**Box 1 Figure 1.**
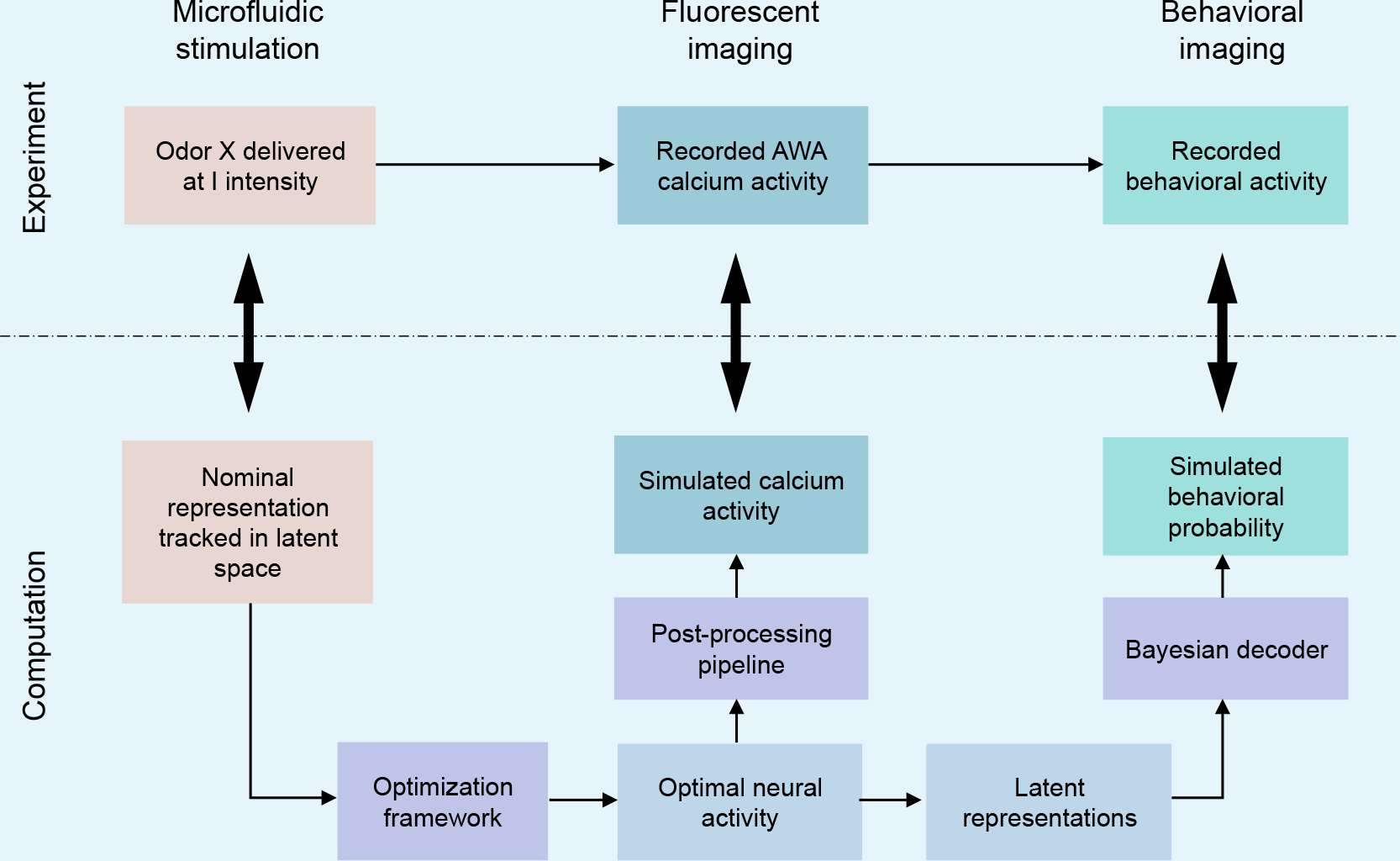
Organization of theoretical and experimental components in our study. Comparisons were made between experimentally observed quantities and simulated variables (marked by bidirectional arrows). The key-components of the computational model are highlighted in purple boxes, while the intermediate computed variables are highlighted in cyan boxes.

#### Optimization framework

The central theoretical premise of this model ***Mallik et al. (2020a***) is the idea that sensory networks through their activity convey information about the peripheral conditions to the downstream regions of the brain. Therefore, if there exists a stimulus specific nominal representation ***z*** in the latent space which embodies information about both the stimulus identity and intensity - then the goal of the sensory units is to create ***v***(*t*) to ‘track’ this representation. More specifically, when there is an odor stimulus, the immediate latent representation must quickly evolve to ***z***. Similarly, when the odor is withdrawn this representation is destroyed and the latent trajectory returns to the origin or the neutral regime of this space.

Under this abstraction, we can posit an objective function that captures the functional significance of sensory neural activity.

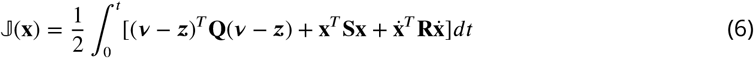

The biological interpretation of this objective function is as follows: the first term penalizes inaccurate latent representation, the second term penalizes the energy expenditure due to neural activity and the third term penalizes rapid fluctuation of neural activity. The optimal neural response strategy obtained by solving the following optimization problem (Eq. 7) ensures that the peripheral stimulus is represented accurately in the latent space while maintaining energy efficiency.

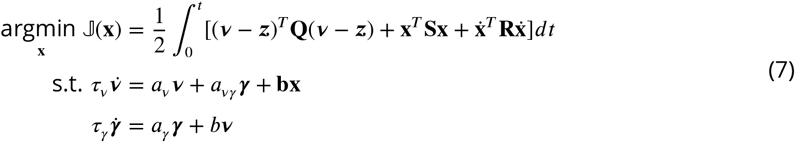

#### Encoding intensity

The mathematical forms for encoding intensity (*c*) under Model 1 and Model 2 are respectively as:

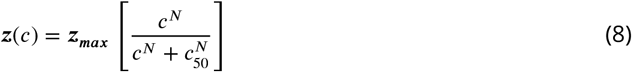

where ***z***_***max***_ is strongest version of the latent representation and *c*_50_ is the stimulus intensity at which the latent representation is at half the strength of its maximum, and,

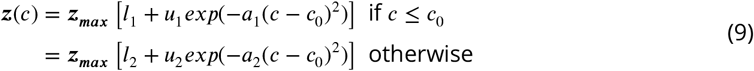

where *c*_0_ is the stimulus intensity at which the latent representation is the strongest. *l*_1_,*u*_1_, and *a*_1_ correspond to the parameters associated with the ascending portion of the psychometric curve, while *l*_2_, *u*_2_, and *a*_2_ are parameters associated with the descending portion of the curve.

The first model of intensity encoding here suggests that with increasing intensity, the system must track a ‘stronger’ latent representation (sigmoidal curve). The second model of intensity encoding indicates that there is ‘preferred’ intensity at which the representation is strongest (a skewed bell curve; see Figure 3).

#### Post-processing steps

The solution of the optimization problem yields instantaneous neural dynamics. To compare neural activity generated by the model with experimentally acquired data from *C. elegans*, we introduced a post-processing pipeline (see Figure 8). This pipeline mirrors the dynamical signal transformations that occur within the AWA neurons of *C. elegans*. The optimization framework generates real-valued activity over time with arbitrary magnitude. The first step in our pipeline therefore was to map the simulated activity to biophysically plausible neural activity patterns. Moreover, GCaMP fluorescence lags calcium dynamics by known on and off time constants ***Larsch et al. (2015***) which introduces latency in the observed response. To model this phenomenon, we passed the time-varying simulated response through a linear system. Borrowing from our prior framework ***Mallik et al. (2020b***), we considered an arbitrary number of neural “units” (20), computed the mean activity across these units, and normalized to pre-stimulation baseline to enable direct comparisons with AWA neuron fluorescence in *C. elegans*.

#### Translating neural codes into behavior

Our goal, fundamentally, is to understand the relationship between measured and modeled chemosensory neural activity and behavior. We begin here with the assumption that the organism displays a finite number of behaviors during our period of observation. Furthermore, there exist certain inherent behavioral dynamics that can be modeled as a Markov process. We assume that the decoded latent variables within *ζ;* form the link between the chemosensory system and behavioral output. Here, we focus on locomotory behavior, specifically forward locomotion, pauses, and reversals. As these relate stochastically to neural activity, the probability of each behavioral state at each time step given the latent information represents model behavior output.

To build this neural-to-latent-to-behavior mapping, we borrowed existing paradigms that have been used to model perceptual decision-making on the basis of sensory evidence. For example, ***Ratcliff (1978***); ***Ditterich (2006***); ***Bogacz et al. (2006***); ***Gold and Shadlen (2007***); ***Mormann et al. (2010***) model decision-making in constrained settings such as ‘interrogation’ or ‘free-response’ ***Bogacz et al. (2006***), wherein a subject must choose between two alternatives within a set amount of time, or on accumulation of a predetermined level of evidence (i.e., a drift-diffusion type integration to bound). Such models can be equivalent to probabilistic inference (Bayesian observer) in many settings ***Bitzer et al. (2014***).

In this work, we model the behavioral outcomes using a stochastic model for evidence accumulation under changing environments ***Veliz-Cuba et al. (2016***). Mathematically, we describe the behavioral output of the model as:

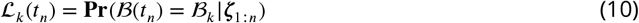

Here, (ℬ_1_, …ℬ) is the finite set of behavior that the organism can execute and *ζ* is the amalgamated latent information from different time-scales (see Figure 9).

**Figure 9.**
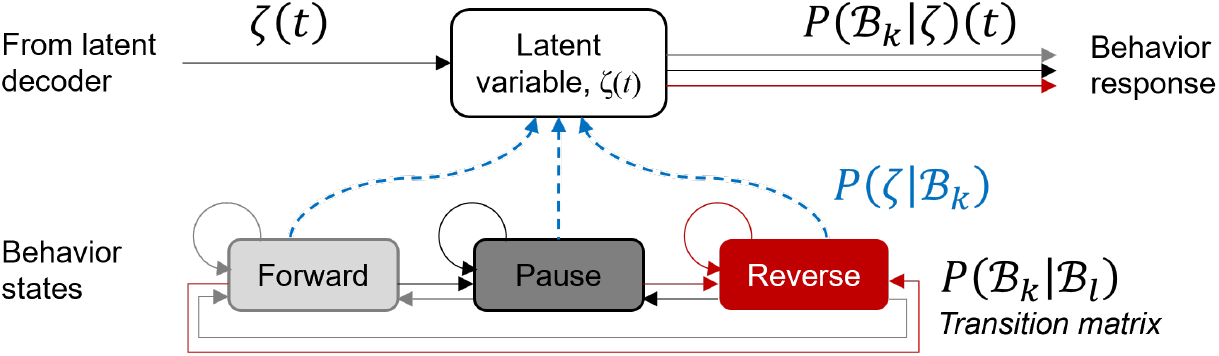
Translating neural codes to behavior

Note that here we discretize time for behavior changes along a coarser timescale than for neural activity. To derive predictions about behavioral output steered by sensory input, we make the following modeling assumptions: (i) Transition of behavior from one state to the other can be modeled via a Markov process. (ii) When the organism is in a specific behavioral state, given the same external stimulation condition persists, the statistics of the input to the decision center remains unchanged. In other words, for every pair of behavioral state and peripheral stimulus condition, there exists a unique random process that characterizes the latent input to the decision center. The information about the stimulus condition and behavioral state is embedded in the mean and covariance, respectively, of these processes. The mathematical details for the model and its parameterization are presented in Appendix 1.

All code to optimize the model and forward predict outcomes was run on a virtualized Microsoft Windows Server 2022 Standard Version 10.0.20348 Build 20348 (PowerEdge R650 System, Intel Xeon Gold 6348 CPU, 512GB RAM) platform, and MATLAB R2023a. Optimization of model parameter differed slightly using other MATLAB versions and computer architectures, but did not significantly affect model predictions.

### Experimental methods

#### *C. elegans* nematode culture

Nematode strains were grown on NGM plates seeded with OP50 bacteria ***Stiernagle (2006***). Strain NZ1091 (*kyIs587 [gpa-6p∷GCaMP2*.*2b; unc-122p∷dsRed]; kyIs5662 [odr-7p∷Chrimson:SL2:mCherry; elt2p∷mCherry]*) expresses the calcium sensor GCaMP2.2b in the AWA chemosensory neurons ***Lagoy (2018***); ***Burnett et al. (2018***). This strain also expresses the light-gated ion channel Chrimson in the same neurons; however, the cofactor required for channel function was omitted for these experiments rendering it nonfunctional ***Larsch et al. (2015***). Young adult *C. elegans* were used in all experiments and synchronized 24 h prior by picking L4 stage worms and transferring them to seeded plates.

#### Fabrication of microfluidic devices for chemical stimulation of *C. elegans*

Microfluidic devices were fabricated according to established methods ***Lagoy et al. (2022***). Briefly, silicon mold masters were fabricated by photolithography using photomasks printed at 25,000 dpi to pattern a 55 - 70 *µ*m layer of SU-8 photoresist. Microfluidic devices were cast to 5 mm thickness by mixing polydimethylsiloxane (PDMS; Sylgard 184, Dow Corning) at a 1:10 ratio, vacuum degassing for 30 min, and curing overnight at 60 ^°^ C. Inlets, outlets, and worm loading ports were punched with a 1 mm diameter dermal punch. Clean devices were sandwiched between glass layers rendered hydrophobic by vapors of tridecafluoro-1,1,2,2-tetrahydrooctyl trichlorosilane (TFOCS; Gelest), with holes drilled through the upper glass slide by a diamond-coated bit, and placed in a metal clamp.

#### Microfluidic device preparation

Microfluidic devices were cleaned, assembled, and degassed in a vacuum desiccator for at least 60 min before experimentation. For behavioral experiments, devices were filled with 5% (w/v) Pluronic F127 through the outlet port to minimize bubble entrapment via its surfactant properties, then flushed with S. basal buffer (100 mM NaCl, 50 mM KPO_4_ buffer, pH 6.0). Neural imaging devices were filled with buffer alone. Reservoirs of loading solutions were prepared as previously described ***White et al. (2023***), purged of bubbles, and connected to the device inlets. A control valve directed buffer or stimulus through the arenas, and flow and fluid switching were monitored prior to loading animals into the arenas.

#### Preparation of chemosensory stimuli

Stimulus fluids were prepared fresh each day prior to experimentation from stock solutions, using analytical grade reagents and ultrapure water (diH_2_O) at room temperature. Animals were maintained during experimentation in 1x S. basal buffer. The attractive odorant diacetyl (2,3-butanedione, Sigma-Aldrich) was serially diluted from a 10^−3^ stock dilution (11.1 mM) in 1x S. basal to final concentrations of 44 nM to 44 *µ*M. To visualize stimulus delivery, fluorescein (25 ng/mL final concentration) was added to diacetyl solutions for neural imaging experiments, and xylene cyanol dye (0.185 mg/mL final concentration) was added for behavior experiments. To prevent motion during neural imaging experiments, animals were paralyzed by addition of the cholinergic agonist tetramisole (1 mM final concentration) ***Larsch et al. (2013***).

#### Fluorescent imaging with chemical stimulation

To quantify neural responses to chemical stimulation, single-plane wide-field, epifluorescent images of multiple *C. elegans* were captured at 10 fps using a Zeiss AxioObserver A1 microscope with a 5x/0.25NA objective, EGFP filter set, and Hamamatsu Orca-Flash 4 sCMOS camera. Animals were injected into the device, preconditioned with five pulses of 15s stimulus (one stimulus each minute), and allowed to paralyze for ∼1 hr in tetramisole buffer before initiation of the stimulus trials. Excitation of GCaMP was achieved by a Lumencor SOLA-LE solid-state lamp triggered by brief (10 ms) pulses synchronized with image capture. An Arduino Nano microcontroller controlled fluidic valves through a ValveLink 8.2 (AutoMate Scientific) controller ***Lawler and Albrecht (2022***); ***White et al. (2023***). Micro-Manager microscope control scripts ***White et al. (2023***) automatically synchronized capture of 24 repeats of a 30s or 55s trial duration with 2-50 second chemical stimulation, beginning at 5s (or 3.5s for 50s duration). Videos were analyzed for neural fluorescence and locomotion using NeuroTracker software in ImageJ, which tracks the position of the neuron over time and integrates background-corrected fluorescent intensity of the neuron soma using an 6 × 6 pixel box ***Larsch et al. (2013***). Using MATLAB, fluorescence intensity *F* (*t*) was normalized by subtracting background fluorescent intensity and then dividing by the initial baseline fluorescence *F*_0_ in the first 3.5 s of each trial before stimulation. Traces from multiple animals were then averaged and reported as Δ*F* /*F*_0_. All software for recording and analyzing neural activity in microfluidic devices are available at *albrechtlab*.*github*.*io*. Peak neural activity was recorded during each stimulation pulse and averaged across the population. A neural Adaptation index was calculated as the mean asymptotic peak neural response (trials 20-24) divided by the maximum peak of the first four trials.

#### Behavioral recording with chemical stimulation

To quantify behavioral responses to chemical stimulation, microfluidic devices containing four 15.5 × 16 mm arenas arranged in a 2 × 2 matrix ***Albrecht and Bargmann (2011***); ***Lawler et al. (2019***) were prepared from masters as described above. Behavioral arenas allowed analysis of 25-30 animals per arena and up to 100-120 animals per experiment. Animal populations were injected into the device, preconditioned with five 15-s stimulus pulses (as for neural imaging experiments) and allowed to acclimate to the microfluidic environment for ≈30 min ***Gray et al. (2005***). Videos were acquired using a 6.6 megapixel PixelLink camera at 30 pixels/mm image resolution and 2 fps using MATLAB recording software. Chemical stimulation timing was syncronized with video recording and controlled by MATLAB via an Arduino Uno and a ValveLink 8.2 (AutoMate Scientific) valve controller. Videos were processed after experimentation using “ArenaWormTracker” MATLAB software to extract behavioral data including position, locomotory speed, and instantaneous behavioral state ***Albrecht and Bargmann (2011***). Short reversals, long reversals, and pirouette turns were combined into a single “Reverse” behavioral state, in addition to “Forward” and “Pause” states. Collisions and unclassified behavior were excluded from analysis.

To translate behavioral probability into quantitative metrics of appetitive versus aversive behavioral responses, we calculated several indices as described in Appendix 2. An instantaneous **Behavior Index** reports population average appetitive behavior for each moment in time, and a behavioral **Response Index** summarizes the mean appetitive response per stimulus trial. For both metrics, positive values indicate attraction, zero indicates no response, and negative values indicate aversion to the stimulus. Finally, a **Habituation Index** compares Response Index magnitude for the average behavioral response index for the last three trials relative to the maximum response index of the first five trials.

#### Statistical analyses

Sample sizes for each experiment are listed in the figure legends. Statistics were performed using one-way ANOVA with Bonferroni’s correction for multiple comparisons or an unpaired two-tailed t test when specified for two sample comparison, using the Statistics and Machine Learning Toolbox in MATLAB. Data are represented as mean ± SEM unless otherwise stated. For curve fitting neural experiment and model data, we used an exponential of the form 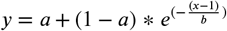, using the Curve Fitting Toolbox in MATLAB.

## Acknowledgments

We thank A. Lammert, A. Philbrook, F. Avery, and V. Kamara for their feedback on the results of this project. Some strains were provided by the CGC, which is funded by National Institutes of Health (NIH) Office of Research Infrastructure Programs (P40 OD010440). D.R.A. was supported by National Science Foundation (NSF) Chemical, Bioengineering, Environmental and Transport Systems Grant 1605679 and Division of Emerging Frontiers Grant 1724026, the NIH Grant R01DC016058, and a Career Award at the Scientific Interface (CASI) from the Burroughs Wellcome Fund.

## Appendix 1

### Outline of the steps to solve the optimization problem

#### Reformulating the optimization problem

The optimization problem as posed in (7) arises from the top-down, normative approach we have adopted for this problem. We denote 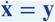. Now the problem can be restated as:

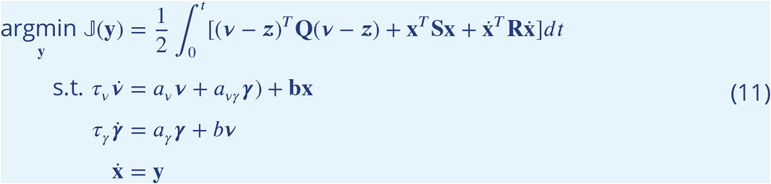

Next, we define an augmented state vector ***ω*** where, ***ω*** = [***γ***^*T*^, ***v***^*T*^, **x**^*T*^, **z**^*T*^]^*T*^. The nominal representation associated with the stimulus i.e., ***z*** remains fixed i.e., ***ż*** = 0 as long as the same peripheral conditions persist. Therefore, we can further reduce (11) to:

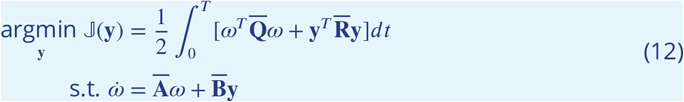

Where,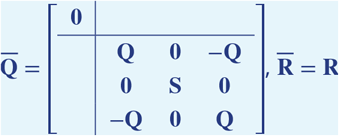,

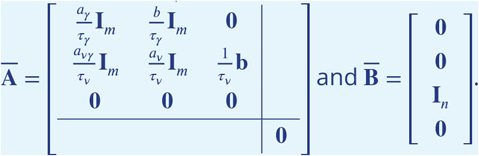

##### Finite Horizon Linear Quadratic Regulator Problem

It turns out that the reduced optimization problem (as outlined in (12)) is a classical control theoretic construct - the Finite Horizon Linear Quadratic Regulator problem ***Anderson and Moore (2007***). A solution to the posited problem (12) exists if the matrix 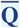 is positive semidefinite and the matrix 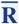 is positive definite. It is straightforward to establish that these conditions are satisfied in our setup. The solution to (12) is therefore given by:

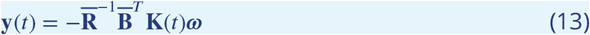

where **K**(*t*) is the solution of the Matrix Riccati Equation:

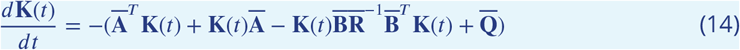

The optimal feedback gain matrix 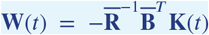 can be decomposed further to obtain the optimal dynamics of the sensory network in a closed form as shown in (2). The detailed steps behind this decomposition can be found in ***Mallik et al. (2020a***). Therefore, **W**(*t*) = [**W** _*γ*_ (*t*): **W**_*v*_ (*t*) : **W**_*x*_(*t*): **W**_*z*_(*t*)].

The connectivity matrices in (2) are given by: 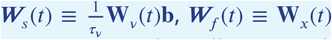 and **r**(*t*) ≡_*z*_(*t*)**z**. Finally, the term ***µ***(*t*) in (2) is obtained by grouping together all terms related to the residual memory ***γ*** (*t*) i.e.,

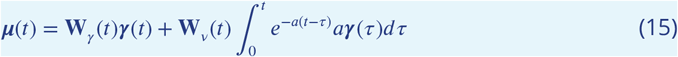

#### Stochastic model of behavior under changing environments

At every timestep, we calculate the likelihood of each behavioral state given the previous and present latent representations. Treating the latent representations as sensory ‘evidence,’ the quantity of interest as expressed in (10) can be reduced as in ***Veliz-Cuba et al. (2016***), resulting in:

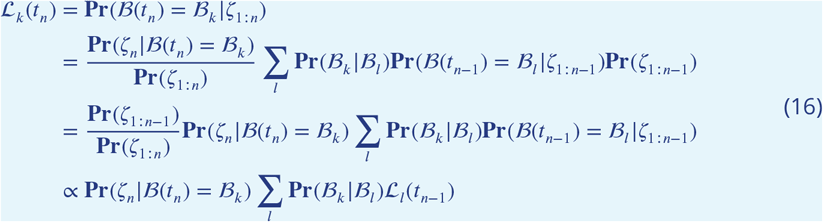

Equation (16) yields that the evolution of behavior over time-steps steered by dynamically encoded sensory information (i.e., here *ζ*) can be modeled via a Hidden Markov Model ***Eddy (2004***). Using Bayes’ theorem we can further write:

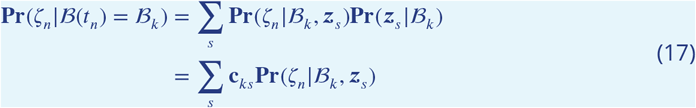

where, *c*_*ks*_ = **Pr**(***z***_*s*_ | ℬ_*k*_) is the probability of a given stimulus condition ***z***_*s*_ (stimulus on vs. off) at that instant of time, given behavior. In the model, this probability will be set as a prior parameter. Plugging (17) in (16), yields

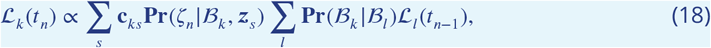

where the HMM architecture is manifested via the transition probabilities **Pr**(ℬ_*k*_ | ℬ_*l*_). Thus,equation (18) indicates that the Bayesian decoder in our construction is fundamentally similar to a downstream decision center that performs computations relevant to behavioral decision making.

### Model parameterization

Below, we provide operational details regarding model construction and parameterization. The model code and experimental data with full parameterization is available at: https://github.com/albrechtLab/Sensory_Memory_Computational_Model.

#### Selecting optimization parameters for generating latent representations and neural responses

In our model simulations, we considered a two-dimensional space, driven by the activity of (hypothetical) neurons that are tuned for either attractive or aversive stimuli, to varying degrees of specificity. To account for variability across organisms, we assumed the existence of 20 such neurons; the nominal modeled AWA neural response is then constructed as the mean of those neurons that favor attractive stimuli.

The penalty matrices in the optimization problem (6) are selected as **Q** = 20𝕀_*m*_, **S** = 2𝕀_*n*_ and **R** = 0.1𝕀_*n*_. The entries in the matrix **b** sample two overlapping Gaussian curves, as described in ***Mallik et al. (2020a***). The parameters *a*_*v*_, *a*_*vγ*_, *a*_*γ*_, *r*_*v*_, *r*_*γ*_ and *b* are selected through numerical optimization. More specifically, we minimize the RMSE error between the simulated Calcium trace and experimentally obtained Calcium trace under constraints evoked from the range of parameters needed to ensure numerical stability of the Matrix Riccati Equation (14). We weight 75% based on the euclidian norm of peak responses (model v. experiment), and another 25% based on the euclidian norm of a point-by-point comparison of the model and experiment dynamics across the whole experiment, with both components normalized by the square-root of vector length. Solving this parameter optimization uses “fmincon” from MATLAB. In our forward simulations, we have used zero mean Gaussian noise with *σ*^2^ = 0.01. Table 1 provides the decoder parameters, reduced to two decimal places, used in our primary simulations, representing mean optimization fits of experimental data from one experiment set at moderate duty cycle (20s (10^−7^ DA) stimulus pulse trials only, 60s ITI).

**Appendix 1 Table 1.**
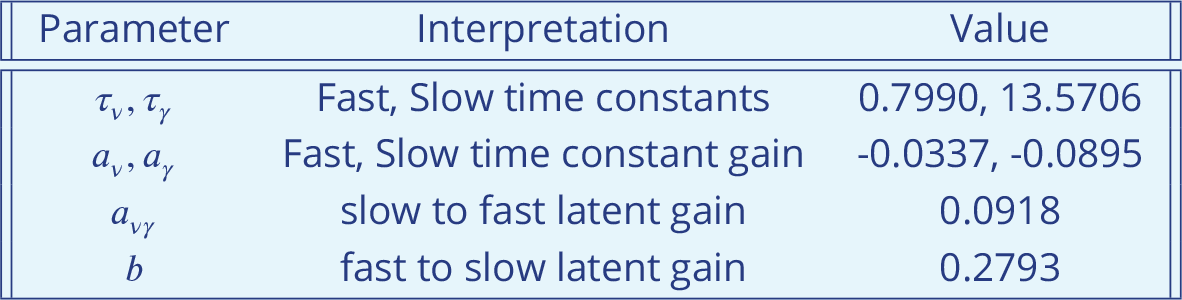
Parameter values for latent decoder and neural response optimization.

#### Selecting parameters for decoding behavior

We previously described a Hidden Markov Model (HMM) architecture to transform latent representations into behavior (18). Within this model, we parameterize the probability of making a latent observation in each behavioral state using a mixture of Gaussian distributions. More specifically,

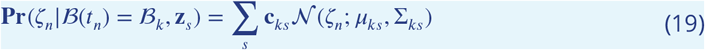

The subscript *s* sums over the conditions of stimulus off (i.e., background buffer *s* = 1) and on (i.e., diacetyl odor *s* = 2). Therefore, here we chose to use a mixture of two multivariate Gaussians to model the emission probability of the HMM. The parameterization of these Gaussian distributions is provided in Table 2.

**Appendix 1 Table 2.**
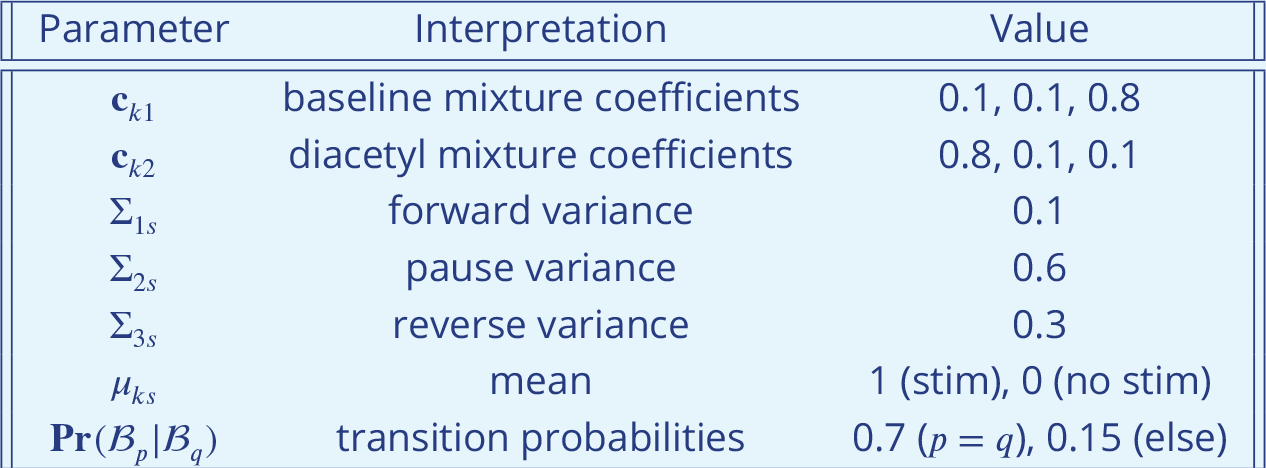
Parameter values for Bayesian behavioral generator.

#### Selecting parameters for concentration

We parameterize the mapping of stimulus intensity or concentration by empirical simulation of habituation behavior across different stimulation timing and concentration followed by comparison with experimental results. From the base optimization concentration, *c* = 1, the target representation *z*_*c*_ could rise or fall, depending on the preferred concentration *c*_0_ of Model 2 (see Figure 1B). Below, different concentration curves generate opposing behavior predictions. As seen in Figure 6, the curve in panel B fits experimental data best.

**Appendix 1 Figure 1.**
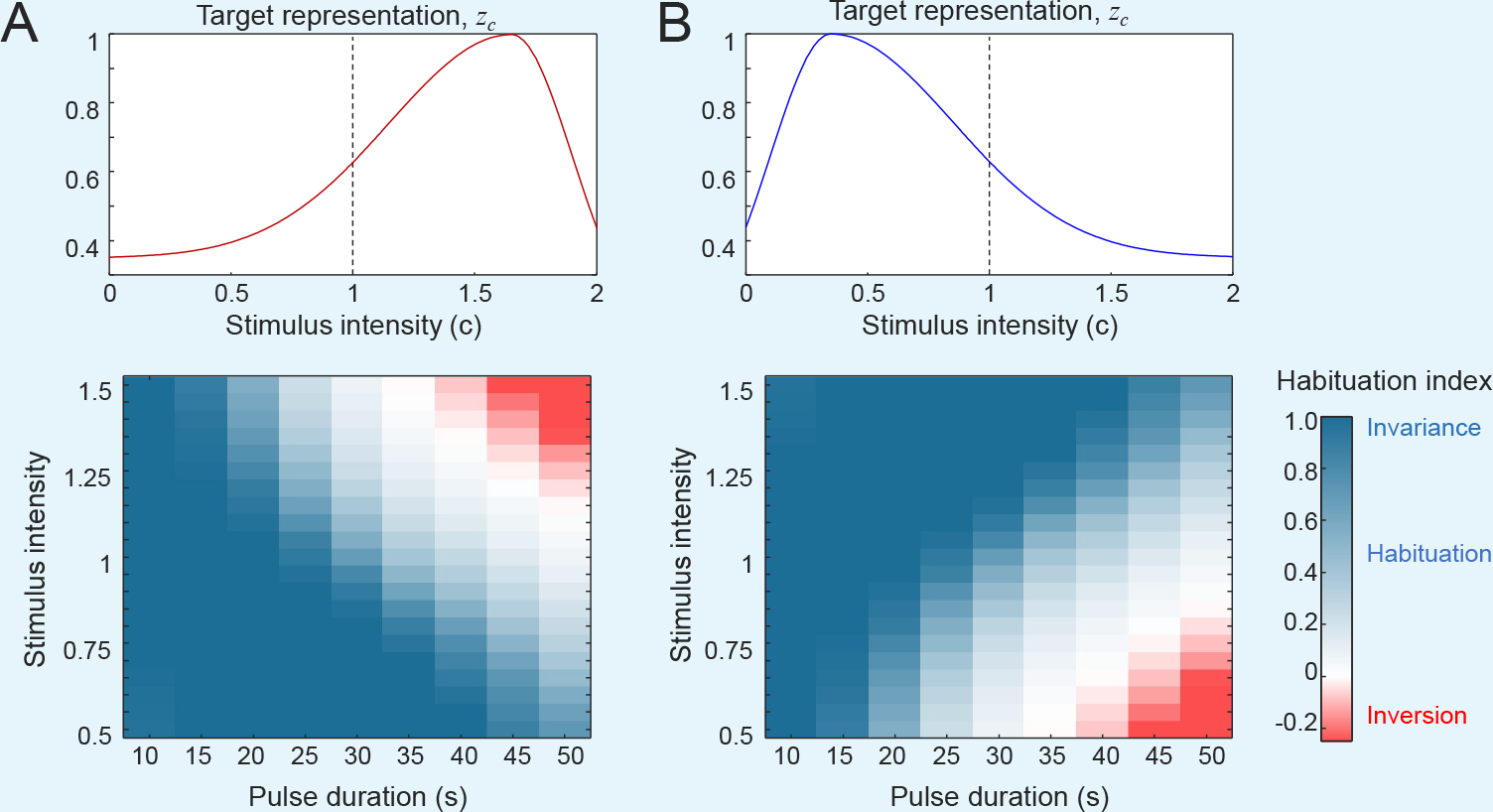
Behavioral ***invariance*** (blue) and ***inversion*** (red) are predicted for long pulse durations and 60 s trial intervals. Different concentration scaling curves change concentrations where inversion is predicted. (**A**) Increasing *z*_*c*_ with stimulus intensity at *c* = 1 predicts inversion at high concentration. (**B**) Decreasing *z*_*c*_ with stimulus intensity at *c* = 1 predicts inversion at low concentration. Experimental data match curve in B (see Figure 6).

## Appendix 2

### Behavior quantification

To quantify animal behavior in response to stimulation, we derived several indices summarizing instantaneous attractive behavior over time (**Behavior Index**), mean attractive response per trial (**Response Index**), and the relative response between the first and last trials (**Habituation Index**). For the Behavior and Response indices, positive values indicate attraction, negative values indicate aversion, and zero indicates no stimulus-evoked change in behavior. For the Habituation Index, a value of 1.0 indicates perfect *Behavioral Invariance* (equal response for first and last trials), zero indicates perfect Habituation (response declines to zero), and negative values represent *Behavioral Inversion* and an opposite response valence.

Experimental recordings of *C. elegans* locomotion yield instantaneous behavioral state probabilities. Behavior tracking software ***Albrecht and Bargmann (2011***) identifies five instantaneous states: Forward, Pause, Short Reversal, Pirouette Reverse, and Pirouette Forward. Here, Short Reversal and both Pirouette states were grouped into a single “Reverse” state representative of avoidance behavior, to match the model output of three behavioral states: Forward, Pause, and Reverse.

As seen in Figures 1F and 4D, attractive behavioral responses are characterized by increased forward probability, decreased reverse probability, and often a slight decrease in pause states upon addition of an attractant stimulus. We derived a single time-varying **Behavior Index** which subtracts the baseline forward probability (outside of stimulus presentation) for each 60-s trial from the forward probability over time (Appendix 2 Figure 1A-D). By resetting the baseline each trial, overall changes in probability are removed (such as changing pause state fractions over time ***Albrecht and Bargmann (2011***); ***Gray et al. (2005***)). Positive Behavior Index represents instantaneous attraction elicited by the stimulus.

To derive a single metric of attractive response to each stimulus trial, we calculated a Behavior **Response Index** representing the difference in Behavior Index during each stimulus (*on1, on2*) and after each stimulus removal (*off1, off2*). A positive Response Index indicates an attractive behavioral response, whereas negative values indicate aversive responses (Appendix 2 Figure 1E,K).

We summarize the degree of behavioral change to repeated stimulation by the **Habituation Index**, or the Response Index at late trials (#24) relative to the first trial (Appendix 2 Figure 1F,L).

**Appendix 2 Figure 1.**
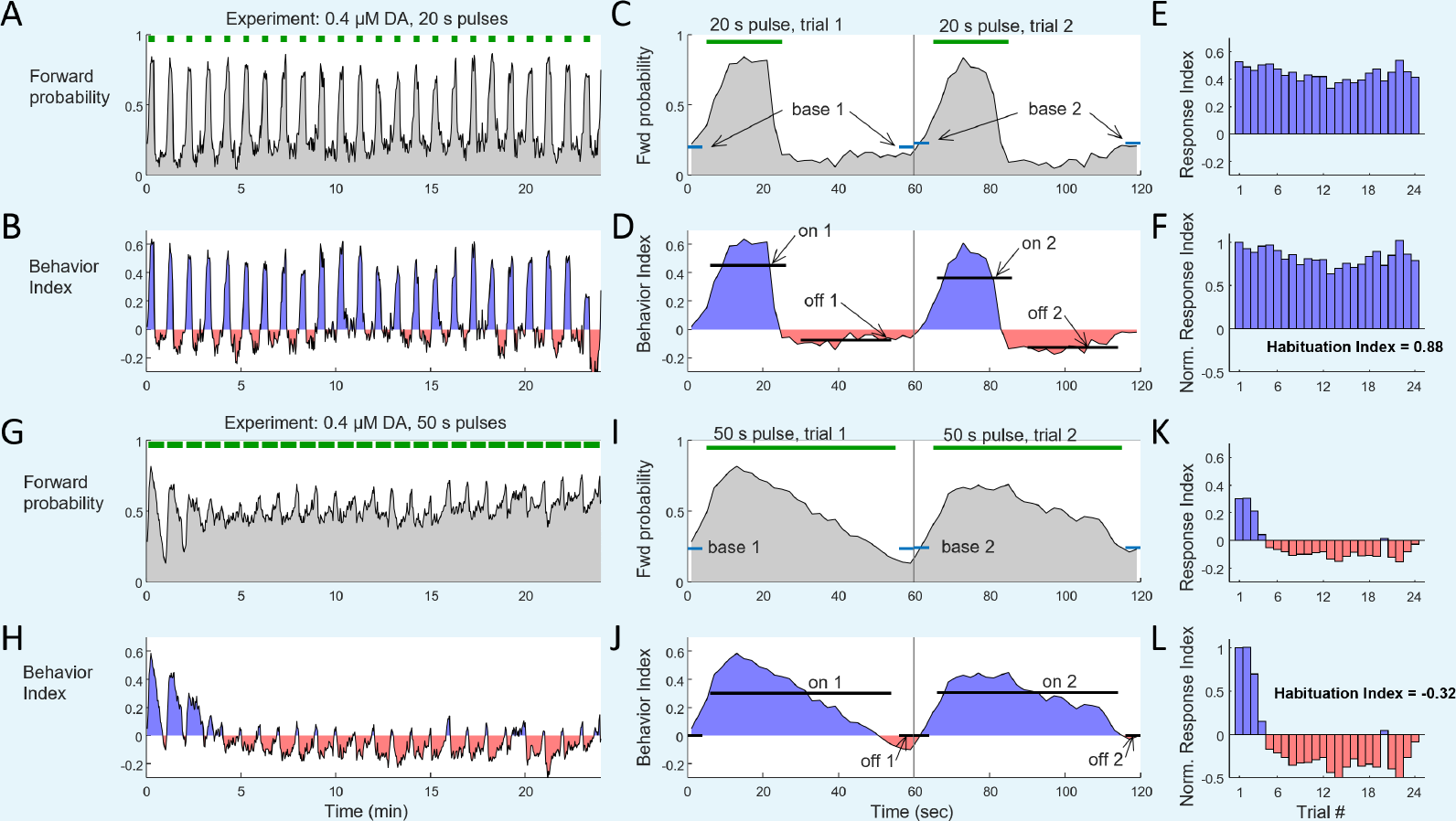
Quantification of behavioral responses. **A**. Forward state probability in response to 20-s pulses of 0.4 uM diacetyl attractive odor. **B**. Behavior Index calculated from Forward probability. **C**. Expansion of first two trials of Forward probability in panel A, indicating measurement of baseline forward probability (base1, base2). **D**. Expansion of first two trials of Behavior Index in panel B, indicating measurement during stimulation (on1, on2) and after stimulation (off1, off2). **E**. Behavior Response Index for all 24 trials, calculated from on - off values in panel D. **F**. Normalized Response Index for all 24 trials calculated relative to trial 1. **G-L**. Same as panels A-F for long 50-s pulses at the same concentration. Note that habituation is minimal for short 20-s pulses (Habituation Index HI = 0.88) and strong for long 50-s pulses, reaching negative HI = -0.32, indicating a change in response valence to aversion at the fifth trial.

